# *Cis*-regulatory mutations co-opting circadian clock regulation underlie naturally selected extreme trait in *Arabidopsis halleri*

**DOI:** 10.1101/2024.10.14.618290

**Authors:** Leonardo Castanedo, Justyna Cebula, Cécile Nouet, Julien Spielmann, Marc Hanikenne, Ute Krämer

## Abstract

*HEAVY METAL ATPase 4* (*HMA4*) is required for the naturally selected traits of zinc/cadmium hyperaccumulation and hypertolerance of in *Arabidopsis halleri*. *Cis*-regulatory alterations and tandem triplication of *AhHMA4* result in substantially elevated transcript levels compared to the closely related non-tolerant non-hyperaccumulator *Arabidopsis thaliana*. Here we identify *cis*-regulatory Metal Hyperaccumulation Elements (MHEs) necessary for *AhHMA4* promoter activity, employing sequence comparisons and motif elicitation analyses combined with progressive deletions and site-directed mutagenesis of promoter-reporter constructs. We report that the promoters of all *AhHMA4* gene copies share a distal MHE1 (consensus TGTAAC), and a proximal pair of MHE2s identical or highly similar to the Evening Element (AAAATATCT). Evening elements are known binding sites of Arabidopsis CIRCADIAN CLOCK-ASSOCIATED 1 (CCA1), a phytochrome-regulated transcription factor in the core circadian clock. We show that the promoter of each *AhHMA4* gene copy, but not of *AtHMA4*, mediates enhanced transcript levels of the reporter and their diel rhythmicity. These functional characteristics are *CCA1*-dependent and recapitulated by a synthetic reporter construct placing the MHE2 pair into the *AtHMA4*-promoter sequence context, according to the example of the *AhHMA4-1* promoter. Consistent with our observations in transgenic reporter lines, *AhHMA4* transcript levels follow a diel rhythm in wild-type *A. halleri* plants. Different from *A. halleri*, we identify complex repressive functionalities co-localizing with an upstream lncRNA and an intron in the 5’ untranslated region of *A. thaliana HMA4*. In summary, our work exemplifies how *cis*-regulatory mutations contributed to the evolution of extreme physiological traits through the co-option of the circadian clock regulatory network.

## Introduction

Evolutionary novelties can result from alterations in coding regions affecting the function, amount or stability of a gene product, or alternatively in non-protein-coding DNA sequences in *cis* that alter the regulation of gene expression (Hill et al. 2021). A growing number of studies support that *cis*-regulatory changes constitute one of the major sources of genetic variation underlying naturally and anthropogenically selected traits in plants and animals (Jacob and Monod 1961; King and Wilson 1975; Stern 1998; Crawford et al. 1999; Wang et al. 1999; Clark et al. 2006; Hanikenne et al. 2008; Hufford et al. 2012; Alonge et al. 2020; Liu et al. 2020; Song et al. 2020). However, studies establishing a mechanistic link between a specific *cis*-regulatory mutation and the corresponding gene expression phenotype remain scarce (Hill et al. 2021; Schmitz et al. 2022). The naturally selected extreme traits of heavy metal hyperaccumulation and hypertolerance are characteristic of the species *Arabidopsis halleri* and absent in closely related species including *Arabidopsis thaliana*. *Cis*-regulatory divergence from the model plant *A. thaliana* at the *HEAVY METAL ATPase 4* (*HMA4*) locus, which encodes a plasma membrane-localized Zn^2+^- and Cd^2+^-exporting pump, is of decisive importance in these extreme traits of *A. halleri* (Hanikenne et al. 2008). The objective of this study was to identify the causal *cis*-regulatory sequence polymorphisms and to obtain information on the *trans* factors involved.

*Arabidopsis halleri* is a diploid stoloniferous perennial and obligate outcrosser within the group of sister species of the intensely studied diploid short-lived selfer *A. thaliana*, from which it diverged between 5 and 10 Mya (Krämer 2010; Novikova et al. 2018). As the only metal hyperaccumulator species within the *Arabidopsis* genus in lineage I of the Brassicaceae, *A. halleri* leaves collected in their natural habitat can contain more than 3,000 (and up to 53,900) µg Zn g^-1^ dry leaf biomass and more than 100 (and up to 3,640) µg g^-1^ Cd, according to the definition of metal hyperaccumulation (Krämer 2010; Stein et al. 2017). Thus, *A. halleri* is capable of accumulating and tolerating more than 10- and up to around 10,000-fold higher metal levels than ordinary plants. In Europe and East Asia, *A. halleri* is among the natural colonizers of so-called calamine metalliferous soils containing high, toxic levels of Zn and Cd from geogenic or anthropogenic sources (Ernst 2006). Species-wide Zn and Cd hypertolerance was confirmed on synthetic media under laboratory conditions (Bert et al. 2003; Becher et al. 2004; Meyer et al. 2010). Different from most of the other > 700 metal hyperaccumulator plant species identified to date, *A. halleri* is a facultative metallophyte (Reeves et al. 2017).

Populations on unpolluted soils containing merely background levels of heavy metals are also hyperaccumulating. Modest Zn hyperaccumulation is also known in the allotetraploid *A. kamchatica*, which arose through the hybridization of *A. halleri* and *A. lyrata* (Paape et al. 2020).

According to cross-species transcriptomics, the transcript levels of tens of metal-related genes are elevated in *A. halleri* compared to *A. thaliana*, mostly constitutively (Becher et al. 2004; Weber et al. 2004; Talke et al. 2006). Subsequently, a reverse genetic approach pinpointed one of these candidate genes, *HMA4*, as the key causal locus making the largest known contribution to both metal hyperaccumulation and metal hypertolerance in *A. halleri* (Hanikenne et al. 2008; Hanikenne et al. 2013). This was supported by the genomic position of *HMA4* within QTL regions mapped for the Zn and Cd hypertolerance traits in a segregating population of a cross of *A. halleri* with *A. lyrata* (Courbot et al. 2007; Willems et al. 2007). By comparison to the closely related non-hyperaccumulating *A. thaliana*, tandem triplication of *HMA4* combined with *cis*-regulatory divergence leading to enhanced promoter strength of all *AhHMA4* gene copies (*AhHMA4-1* to *AhHMA4-3*), resulted in 6- to 50-fold elevated *HMA4* transcript levels in *A. halleri* (Talke et al. 2006; Hanikenne et al. 2008; Hanikenne et al. 2013).

Beyond enhanced *HMA4* gene product dosage (Hanikenne et al. 2013), the evidence available to date does not support any predominant role of divergent transcript localization or divergent functions of the encoded proteins among the *AhHMA4-1* to *AhHMA4-3* gene copies, or compared to *AtHMA4* (Krämer 2010; Nouet et al. 2015). Ectopic Gene Conversion, also addressed as Inter-locus Gene Conversion, among *AhHMA4-1*, *AhHMA4-2* and *AhHMA4-3* is well-documented and reflected by ≥ 99% coding sequence identity between gene copies, further supporting that their protein-coding sequences undergo concerted evolution (Hanikenne et al. 2008; Hanikenne et al. 2013). Nucleotide polymorphism is consistent with positive selection and a hard selective sweep in the genomic region comprising *AhHMA4-1* to *AhHMA4-3* (Hanikenne et al. 2013).

Here we identify the divergent *cis*-regulatory sequences required for elevated activity of the promoters of the *HMA4* gene copies of *A. halleri* compared to *A. thaliana HMA4.* We employ sequence comparisons and analyses, as well as promoter deletion series, promoter mutation and segmental promoter swap constructs. In the promoters of all three *AhHMA4* gene copies, we identify the conserved *cis*-regulatory enhancer elements Metal Hyperaccumulation Element 1 and 2 (MHE1 and MHE2), which contribute to high transcript levels of a downstream reporter gene in the *A. thaliana* genetic background and are absent in the *A. thaliana HMA4* promoter. By contrast, we report that *A. thaliana HMA4* is targeted by complex repressive *cis*-regulatory functionalities, which co-localize with a distal upstream lncRNA, an intron region corresponding to the 5’-untranslated region of the transcript. MHE2 sequences are identical or highly similar to the known evening element, a known binding site for the MYB family transcription factor CIRCADIAN CLOCK ASSOCIATED 1 (CCA1) that mediates light-dependent regulation and forms part of the core oscillator of the Arabidopsis circadian clock (Wang et al. 1997; Wang and Tobin 1998; Alabadí et al. 2002). We show that both elevated levels and diel dynamics of reporter gene transcript levels depend on the combination of the functionalities of *CCA1* in *trans* and MHE2 in *cis*. Introducing both copies of MHE2 from the *AhHMA4-1* promoter into an *AtHMA4* promoter context recapitulates strongly elevated reporter gene transcript levels and their diel rhythms. Endogenous *HMA4* transcript levels exhibit similar diel dynamics in *A. halleri*, thus supporting the validity and relevance of our findings. Taken together, our work constitutes a major step forward in our understanding of the mechanistic basis of an extreme trait syndrome in plants and provides an example of how *cis*-regulatory divergence contributed to the evolution of an extreme physiological trait through the co-option of a core circadian clock transcriptional regulator.

## Results

### Identification of regions governing functional divergence between the *A. halleri HMA4-1* and the *A. thaliana HMA4* promoter

Previous research demonstrated that metal hyperaccumulation and the full extent of metal hypertolerance in *A. halleri* depend on strongly elevated *HMA4* transcript levels (Talke et al. 2006; Courbot et al. 2007; Hanikenne et al. 2008). This was attributed to *cis*-acting sequence differences between *A. halleri* and *A. thaliana HMA4* by using constructs of genomic DNA segments fused to the *β-GLUCURONIDASE* (*GUS*) reporter gene in the genetic backgrounds of both species, *A. halleri* and *A. thaliana* (Hanikenne et al. 2008). In the present study, we aimed to identify the specific *cis*-regulatory sequence alterations in *A. halleri HMA4* genes that confer strongly enhanced GUS reporter activities by comparison to the corresponding regions of *A. thaliana HMA4* which mediate substantially lower GUS activities (Hanikenne et al. 2008). For this purpose, we generated series of promoter deletion constructs fused to a *GUS* reporter gene, and we introduced these into *A. thaliana*, given that *HMA4* promoter activities are independent of the *A. halleri* or *A. thaliana* genetic backgrounds (Hanikenne et al. 2008). Although generally feasible, transgenic approaches in *A. halleri* remain highly time-consuming and laborious to date. Among the promoter regions of the three *HMA4* gene copies of *A. halleri* (accession Lan3.1), *AhHMA4-1*, *AhHMA4-2* and *AhHMA4-3*, the sequence of *AhHMA4-1_P_* is the most similar to *AtHMA4_P_* (accession Col-0), which facilitated direct comparisons (Supplementary Fig. S1 A-D) (Hanikenne et al. 2008; Hanikenne et al. 2013).

Our full-length *AhHMA4-1_P_* construct comprised a genomic fragment of 2,326 bp in length, consisting of 1,602 bp upstream of the transcriptional start site, the 5’ UTR including exon 1, an intron, and the beginnning of exon 2, followed by the initial 30 bp of the *AhHMA4-1* coding sequence (Hanikenne et al. 2008). We designed progressive 5’ deletions, with breakpoints guided by the boundaries of segments exhibiting sequence similarity between the promoters of *A. halleri HMA4-1* and *A. thaliana HMA4* (Fig. 1A, Supplementary Figure S1D). Deletion of the initial 904 bp at the 5’-end of *AhHMA4-1_P_*, addressed here as the distal region (DR), to generate the *AhHMA4-1_P_* Δ698 construct, caused a strong decrease in both *GUS* transcript levels and specific GUS enzyme activities of total protein extracts down to residual levels of 37%, 24%, and 35% on average, in shoots, roots, and whole seedlings, respectively (Fig. 1A, “Enhancing Region 1”; Fig. 1B and C, compare full-length (FL) *AhHMA4-*1*_P_* FL with Δ698; Suppl. Fig. S1E and F, compare *Ah-1_P_* FL with ΔDR). Additional progressive deletions of the downstream intermediate region (IR) of 568 bp in length had no or only minor effects (Fig. 1A-C, IR, from Δ438 to Δ130). Deletion of the intron in the *AhHMA4-1_P_* construct had no significant effects by comparison to either the full-length promoter or the truncated promoter lacking both DR and IR (Fig. 1B and C, compare FL with FLΔi and Δ130 and Δ130Δi, and Suppl. Fig. S1E and F, compare ΔDR with ΔDR to ΔDIRΔi). The additional deletion of the 44-bp-long proximal region (PR) to generate the *AhHMA4*-*1_P_* Δ86Δi construct led to decreases in *GUS* transcript levels down to 2%, 3%, and 1%, of the full-length promoter in seedlings, shoots and roots, respectively (Fig. 1A, “Enhancing Region 2”; Fig. 1B, Δ86Δi; Suppl. Fig. S1E, ΔDIPRΔi). In these lines, residual *GUS* transcript levels and specific GUS activities were very low and indistinguishable from control transformants devoid of a promoter upstream of the *GUS* reporter gene (Fig. 1B and C, compare Δ86Δi and ev, Suppl. Fig. 1E and F).

**Figure 1.**
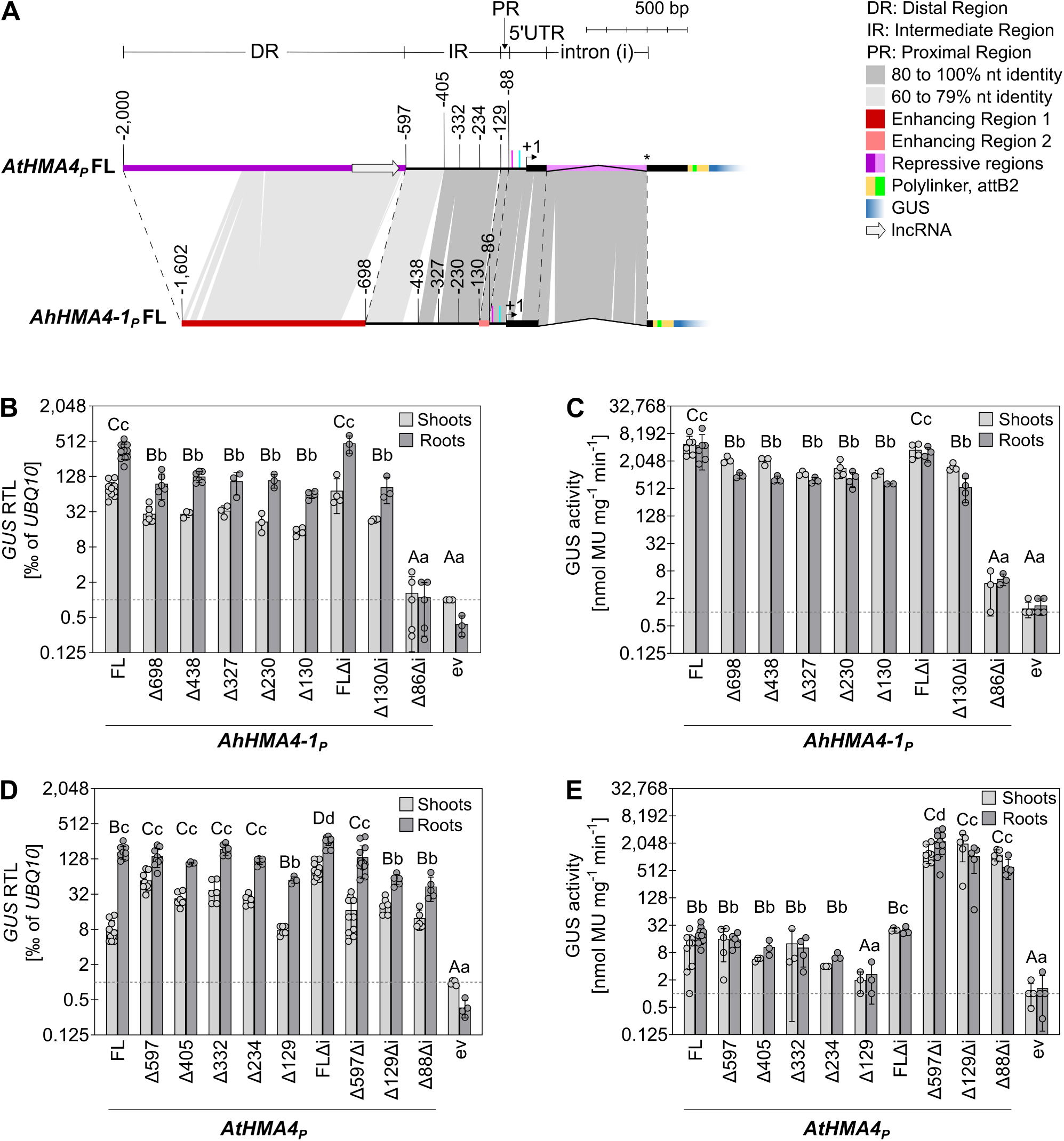
Analysis of deletion series of *AtHMA4* and *AhHMA4-1* promoter regions in transgenic reporter lines of *A. thaliana*. **(A)** *GUS* reporter constructs for *AtHMA4_P_* and *AhHMA4-1_P_*. Numbers indicate distances in bp from the transcriptional start site (+1) for each of the breakpoints implemented in the promoter deletion series of reporter constructs introduced into *A. thaliana*. Positions of putative CAAT/TATA boxes are marked by magenta/cyan vertical lines. 5’UTR, 5’ untranslated region; *,translational start site (vertically aligned between promoters); FL, full-length. **(B-E)** Relative *GUS* transcript levels (B and D) and specific GUS enzyme activities (C and E) for full-length *AhHMA4-1_P_* (B and C), *AtHMA4_P_* (D and E), and their respective deletion series in transgenic *A. thaliana GUS* reporter lines. Bars show mean ± SD (*n* = 3 to 14) of independent transgenic lines, with each datapoint representing the mean of three replicate multi-well plates of qPCR reactions (B, D) or enzyme assays (C, E). Data are from one representative experiment out of a total of two independent experiments. Different characters indicate statistically significant differences based on one-way non-parametric ANOVA with Dunńs multiple comparison test (*P* < 0.05). Δi, deletion of the intron (see A); ev, transformants with a construct lacking any promoter fragment upstream of the *GUS* gene (see Materials and Methods); MU, 4-methylumbelliferone.

The 2,799-bp-long full-length promoter fragment of *A. thaliana HMA4* included 2,000 nucleotides upstream of the transcriptional start site (Hanikenne et al. 2008). *GUS* transcript levels driven by *AtHMA4_P_* were 14% in seedlings, 8% in shoots, and 54% in roots, of *GUS* transcript levels in the lines carrying full-length *A. halleri HMA4* promoters (Fig. 1B and D, see also Fig. 2P). GUS enzyme activities directed by *AtHMA4_P_* were far below those observed in the *AhHMA4_P_* lines and amounted to less than 0.5% (Fig. 1C and E, see also Fig. 2Q), in agreement with previously published results (Hanikenne et al. 2008). In relation to *GUS* transcript levels, specific GUS reporter enzyme activities in full-length promoter-reporter lines were considerably lower for *AtHMA4_P_* than for *AhHMA4-1_P_* (Fig. 1B-E, Suppl. Fig. S2A-D, *AhHMA4-1_P_*FL is among constructs showing slopes of 8.2 in roots and 65 in shoots, contrasting with *AtHMA4_P_*FL among constructs with slopes of 0.09 in roots and 0.07 in shoots). Thus, GUS enzyme activities were in qualitative agreement with *GUS* transcript levels for *AhHMA4*-*1_P_*, but not for *AtHMA4_P_*. Compared with the full-length *AtHMA4_P_* construct, the combined deletion of both the DR and the intron resulted in about 100-fold increased GUS enzyme activities in total protein extracts of both shoots and roots (Fig. 1E, compare Δ597Δi to FL). In these *AtHMA4_P_* Δ597Δi lines, GUS activity expressed as a function of *GUS* transcript levels reached similar magnitudes as in full-length *AhHMA4-1_P_* lines (Fig. 1D and E, Suppl. Fig. S2B and D, *AtHMA4_P_* ΔDRΔi among constructs showing slopes of 17 in roots and 84 in shoots. This suggested that both the DR and the intron of *AtHMA4_P_* were independently able to strongly repress GUS reporter activity, because the deletion of either the DR or the intron individually had no effect on GUS activity, and the deletion of both the DR and the intron in combination was required for increased GUS activity. At the *GUS* transcript level, deletion of the DR of *AtHMA4_P_* resulted in a 7-fold increase in shoots and no significant change in roots (Fig. 1D, compare FL and Δ597). Moreover, deletion of the intron of full-length *AtHMA4_P_* caused an 11-fold increase in *GUS* transcript levels in shoots and a statistically significant 2-fold increase in roots (Fig. 1A and 1D, compare FL and FLΔi). Consequently, the DR and the intron of *AtHMA4_P_* each appeared to have repressive effects on *GUS* transcript levels primarily in shoots. Yet, these effects appeared to be complex, because the simultaneous deletion of both the DR and the intron resulted in *GUS* transcript levels at similar magnitudes as observed for the full-length *AtHMA4_P_* construct. Furthermore, our results suggested that the region of *AtHMA4_P_* between 234 and 129 nucleotides upstream of the transcriptional start site had a 2- to 3-fold enhancing effect on *GUS* transcript levels in the presence of the intron (Fig. 1D, compare Δ234 and Δ129), consistent with similar changes in GUS activity (Fig. 1E). In the absence of the intron, we detected a similar effect only in the roots. In contrast to *AhHMA4-1_P_*, deletion of the PR in *AtHMA4_P_* did not cause any change in *GU*S transcript levels or GUS activity (Fig. 1D and E, compare Δ129Δi with Δ88Δi). Taken together, our results are consistent with the presence of *cis*-regulatory elements causing enhanced transcription located in the DR (“Enhancing Region 1”, between - 1602 and −698) and close to the transcriptional start site in the PR (“Enhancing Region 2”, between −130 and −86) of the *A. halleri HMA4-1* promoter. For *AtHMA4*, the analysis of our reporter constructs suggested the presence of complex repressive functionalities involving the DR and an intron in the 5’ UTR. These functionalities had by far the strongest effects at the protein level. In brief, our results implicate two enhancing regions in the *A. halleri HMA4-1* promoter and two repressive regions upstream of *AtHMA4*.

**Figure 2.**
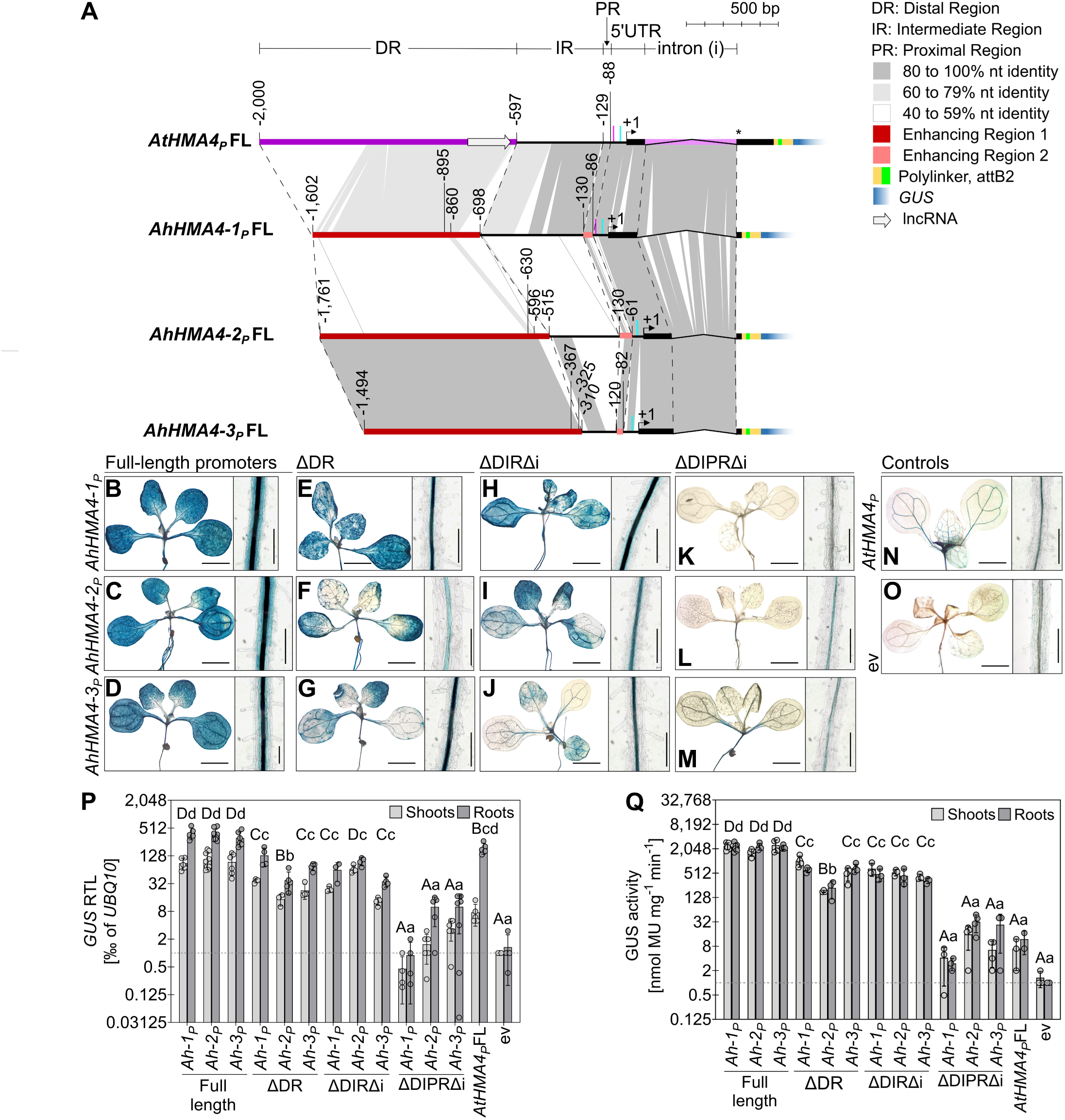
Identification of two enhancing regions in homologous positions of the promoters of all three *A. halleri HMA4* gene copies. (A) *GUS* reporter constructs for *AtHMA4*, *AhHMA4-1*, *AhHMA4-2*, and *AhHMA4-3* promoters. See also Fig. 1A. **(B-O)** Histochemical detection of GUS activity in rosettes (left) and the root hair zone of roots (right) of representative transgenic *A. thaliana GUS* reporter lines. Size bars: 2 mm for rosettes, 0.2 mm for roots. **(P and Q)** Relative *GUS* transcript levels and specific GUS enzyme activities in transgenic *A. thaliana* reporter lines. Bars show the mean ± SD (*n* = 3 to 7) of independent transgenic lines, with each datapoint representing the mean of three technical replicates (see Fig. 1). *Ah-1*, *AhHMA4-1*; *Ah-2*, *AhHMA4-2*; *Ah-3*, *AhHMA4-3*; ΔDIR, deletion of both the distal and the intermediate regions, ΔDIPR, deletion of the distal, intermediate, and proximal regions (see also Fig. 1B-E).

### All three *AhHMA4* gene copies share two homologous enhancing *cis*-regulatory regions

Because the promoters of all three paralogous *AhHMA4* gene copies of *A. halleri* are known to confer high levels of specific GUS activity in total protein extracts (Hanikenne et al. 2008), we hypothesized that “Enhancing Region 1” and “Enhancing Region 2” of *AhHMA4-1_P_* are functionally conserved in both *AhHMA4-2_P_* and *AhHMA4-3_P_* (Fig. 2A). The full-length *AhHMA4-3_P_* construct (total length 2,060 bp) comprised 1,494 bp upstream of the transcriptional start site (Hanikenne et al. 2008; Nouet et al. 2015). By comparison, deletion of the DR (from 1,494 to −310), which was delineated based on synteny with *AhHMA4-1*, caused reductions in both *GUS* transcript levels and specifiy GUS activity down to 30% in shoots and seedlings, and 25% in roots, very similar to our observations for *AhHMA4-1_P_* (Fig. 2B, D, E, G, P and Q, and Suppl. Fig. S1E and F, compare FL with ΔDR for *Ah-1_P_* and *Ah-3_P_*). For *AhHMA4-2_P_* (total length 2,293 bp), which included 1,761 bp upstream of the transcriptional start site, deletion of DR (from - 1,761 to −515) resulted in only 10% remaining promoter activity according to both *GUS* transcript levels and GUS activity and irrespective of the analyzed tissue types (Fig. 2C, F, P, and Q, and Suppl. Fig. S1E and F, compare full-length with ΔDR for *Ah-2_P_*). Similar to what we observed for *AhHMA4-1_P_*, the additional deletion of PR in constructs lacking DR, IR, and the intron, resulted in very strongly decreased activities of *AhHMA4-2_P_* and *-3_P_* (Fig. 2H-J, K-M, N-Q, and Suppl. Fig. S1E and F, compare ΔDIRΔi with ΔDIPRΔi for *Ah-1_P_*, *Ah-2_P_*, and *Ah-3_P_*). This suggested that both “Enhancing Region 1” and “Enhancing Region 2” are functionally conserved in the promoters of all three *HMA4* gene copies of *A. halleri*. Besides this, a small difference was observed between *AhHMA4-2_P_* and the promoters of the other two *A. halleri HMA4* gene copies. Deletion of both IR and the intron in *AhHMA4*-*2_P_* lacking DR led to a small increase in *GUS* transcript levels up to 3-, 4-, and 2-fold higher levels in seedlings, shoots, and roots, respectively, which was also partly apparent at the level of GUS activity (Fig. 2F, I, P, and Q, and Suppl. Fig. S1E and F, compare ΔDR with ΔDIRΔi for *Ah*-*2_P_*). Thus, either the IR of the promoter region, or the intron in the 5’-UTR, of *AhHMA4-2* may uniquely harbour a moderately repressive element, which we did not analyze further here.

### Identification of *cis*-regulatory elements in the *A. halleri HMA4* promoters

Further dissecting Enhancing Region 1 (ER1) present in *A. halleri HMA4* promoters using the next series of deletion constructs of *AhHMA4-1_P_* identified a predominant contribution by the segment between positions −909 and −806 of the DR of *AhHMA4-1_P_* (Fig. 3A, see Suppl. Figs. S3 and S4A). The deletion of this promoter segment resulted in a decrease in both *GUS* transcript levels and specific GUS activity down to a residual level of about 30% and 37% in shoots and seedlings, and 25% in roots (Fig. 3D-G, P, and Q, and Suppl. Fig. S4C-J). Within this segment, we identified an approximately 36-bp-long region showing an elevated degree of sequence similarity among the *A. halleri HMA4* promoters and of shared divergence from *AtHMA4_P_* (ER1^+^, −895 to −860 in *AhHMA4-1_P_*, Fig. 3B, Suppl. Figs. S1D, S3 and S4B). By using multiple sequence alignments and motif elicitation analysis across ER1^+^ of the promoters of all three *AhHMA4* gene copies from a diverse set of *A. halleri* populations, we identified a conserved putative *cis*-regulatory element of 12 bp in length (−895 CTT TGT AAC CAT - 884) (Suppl. Fig. 5a, Suppl. Dataset S1). This element contains the 7-bp core motif TGTAACC (−892 to −886), which is not present in *A. thaliana* (designated Metal Hyperaccumulation Element 1, MHE1; Suppl. Fig. S4B, see Methods). We designated this element MHE1a, because a second copy of the MHE1 core motif, MHE1b, appears to be present upstream at positions −914 to −908 of *AhHMA4-1_P_* (Fig. 3B, and Suppl. Figs. S3 and S4B), but not in *AhHMA4-2_P_* and *AhHMA4-3_P_*. Indeed, disruption of MHE1b in the Δ909 construct led to a subtle, statistically significant decrease in promoter activity at the *GUS* transcript level in roots (Suppl. Fig. S4I, compare Δ1,362 and Δ909), which we did not detect based on GUS activity or in whole seedlings, however (Fig. 3P and Q, Suppl. Fig. S4J). The *AtHMA4_P_* sequence −684-TGTAATC-678 corresponding to MHE1b of *AhHMA4-1* was highly similar, with a single C-to-T exchange at the penultimate position, whereas MHE1a was absent in *AtHMA4_P_* (Fig. 3B, Suppl. Figs. S3, S4B, and S5A).

**Figure 3.**
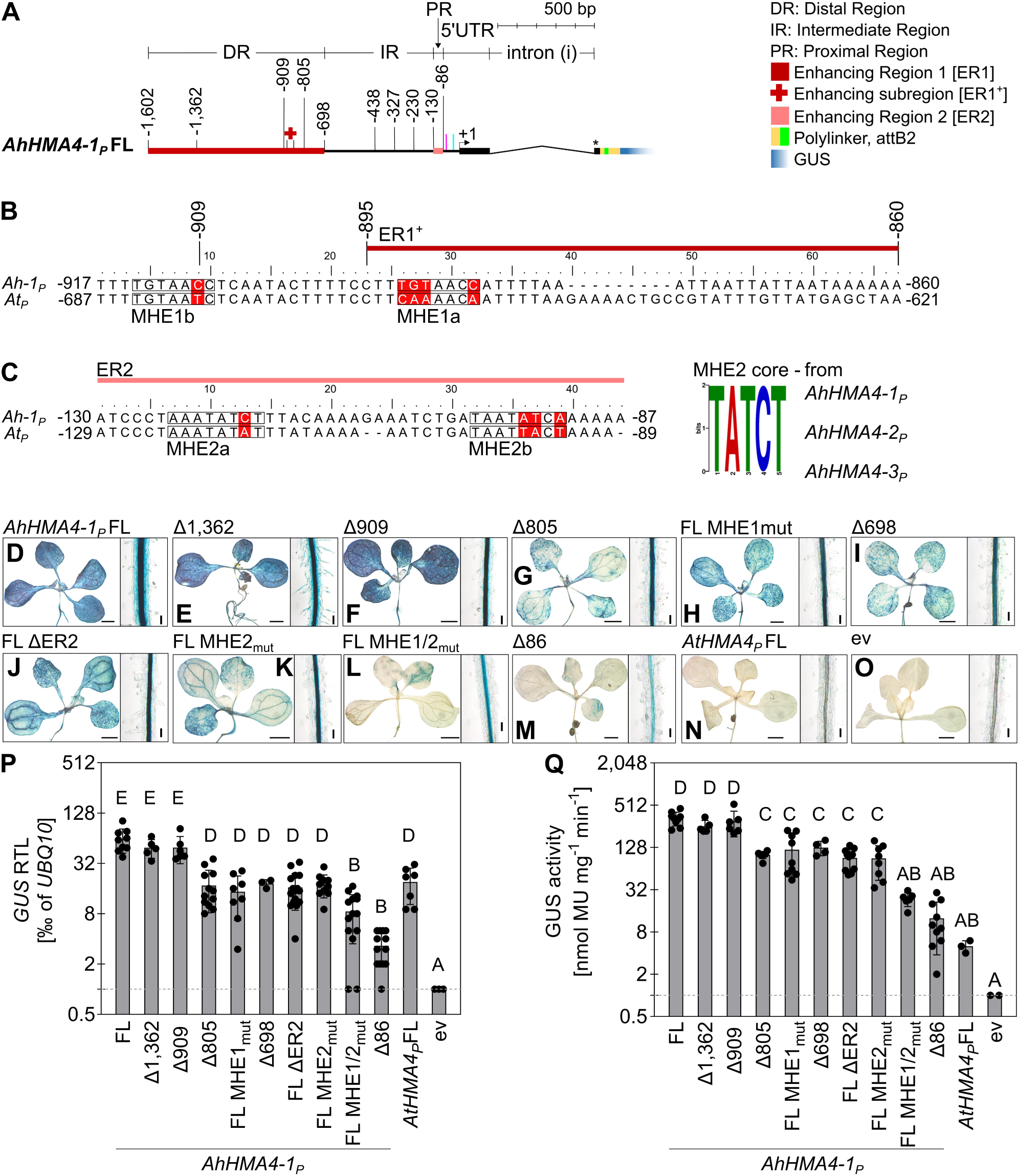
*Cis*-regulatory enhancer elements MHE1 and MHE2 are required for activity of the *AhHMA4-1* promoter. **(A-C)** Scheme of *AhHMA4-1_P_* constructs (A), as well as alignments of the ER1^+^ region (B) and the ER2 region (C) of *AhHMA4-1_P_* with the homologous regions of *AtHMA4_P_* (left) and core MHE2 motif (right; see Suppl. Fig. S6). The sequences of the putative *cis*-regulatory elements in the effective subregion of ER1, ER1^+^ (red cross symbol in A, red bar in B), are boxed: Metal Hyperaccumulation Element 1a (MHE1a) and a second identical copy upstream (MHE1b, B), as well as MHE2a and MHE2b in ER2 (light red bar, C). Between-species differences are shown on a red background, and they also indicate the mutations introduced into *AhHMA4-1_P_* (conversion to the corresponding *AtHMA4_P_* sequence) to disrupt MHE1a/b and/or MHE2a/b (B and C). See also Fig. 1A. The sequence logo of the core MHE2 motif shared by the promoters of all three AhHMA4 gene copies in *A. halleri* is shown. **(D-O)** Histochemical detection of GUS activity in rosettes (left) and the root hair zone of roots (right) of representative transgenic *A. thaliana GUS* reporter lines. Size bars: 1 mm (rosettes), 0.2 mm (roots). **(P and Q)** Relative *GUS* transcript levels (P) and specific GUS enzyme activities (Q) in whole seedlings of transgenic *A. thaliana GUS* reporter lines. Bars show mean ±SD (*n* = 3 to 15) independent transgenic lines. Each datapoint represents the mean of three technical replicates (see Fig. 1). Different characters indicate statistically significant differences based on one-way non-parametric ANOVA, followed by a Dunn’s multiple comparison test (*P* < 0.05). ΔER2, deletion of ER2 (see C) in a full-length promoter; mut, mutated (see B, C); MHE1/2_mut_, MHE1 and MHE2 mutated. See also Fig. 1.

A multiple sequence alignment of the PR, which corresponds to Enhancing Region 2 (ER2) of promoters of the three *A. halleri HMA4* gene copies, indicated sequence similarities among only small segments therein (Fig. 3C, Suppl. Fig. S6A). We identified two copies of an 8-bp-long motif −124-CACTATCT-117 nt (Suppl. Fig. S6A, Suppl. Dataset S2), with the core motif TATC(T/A), which we designated MHE2 (Metal Hyperaccumulation Element 2) (Fig. 3C). None of these two copies of MHE2 were present in the PR of *AtHMA4_P_*, and only one of them, MHE2a, was present in in *AlHMA4_P_* (Suppl. Fig. S6A). In conclusion, MHE1a, MHE2a and MHE2b were present in the promoters of all three *AhHMA4* gene copies (see Suppl. Figs. S3, S5A and S6A). Next we tested whether the putative *cis*-regulatory elements MHE1 and MHE2 of *AhHMA4-1_P_* are necessary for its promoter activity. We replaced the characteristic nucleotides by converting MHE1 and MHE2 into the sequences of the corresponding positions of *AtHMA4_P_* using site-directed mutagenesis (Fig. 3A-C). Compared to full-length wild-type *AhHMA4-1_P_*, a mutated *AhHMA4-1_P_* carrying five single-nucleotide replacements to alter both copies of the MHE1 motif conferred only about 25% residual GUS activity (Fig. 3D, H, P and Q, compare MHE1_mut_ with FL for *AhHMA4-1_P_*). Residual GUS activity of the mutated *AhHMA4-1_P_* was comparable to that for constructs in which the MHE1-containing region or the entire DR was deleted (Fig. 3G to I, P and Q, compare MHE1_mut_ with Δ805 and Δ698). Similarly, upon converting the two MHE2 motifs of full-length *AhHMA4-1_P_* into the corresponding sequence of *AtHMA4_P_* through four single-nucleotide replacements, we observed only about 25% residual GUS activity, indistiguishable from the residual activity observed for a construct in which only the PR corresponding to ER2 was deleted in an otherwise full-length *AhHMA4-1_P_* Fig. 3C and D, J-K, P and Q, compare MHE2_mut_ with FL and FL ΔER2). Site-directed mutagenesis of both MHE1 and MHE2 in combination led to a substantial reduction in GUS activity down to only about 10% of that observed for the full-length *AhHMA4*-*1_P_* construct (Fig. 3B-D, L, M, P, and Q, compare MHE1/2_mut_ with Δ86 and FL). In summary, these results supported the hypothesis that the two MHE1 motifs in the DR and the two MHE2 motifs in the PR comprise *cis*-regulatory elements that are necessary for the full, strongly elevated promoter activity of *AhHMA4-1_P_*, by comparison to *AtHMA4_P_*.

We also examined whether MHE1 and MHE2 of *AhHMA4-1_P_*, when introduced into *AtHMA4_P_*, are sufficient to confer enhanced promoter strength to *AtHMA4_P_*. We used site-directed mutagenesis to introduce MHE1 and MHE2 of *AhHMA4-1_P_* into *AtHMA4_P_* Δ754Δi, which lacks the intron and comprises a shortened DR of only 157 bp of its 3’ end (Fig. 4A). *AtHMA4_P_* Δ754Δi comprises both the sequence segment (−665 to −621) of *AtHMA4_P_* showing microsynteny to ER1^+^ in *AhHMA4-1_P_* (−895 to −860; see Fig. 3B) and the PR corresponding to ER2 in *AhHMA4-1_P_* (Fig. 3C). Functionally, *AtHMA4_P_* Δ754Δi was equivalent to full-length *AtHMA4_P_* with respect to GUS transcript levels and GUS activity (Fig. 4B, G, L and M). For a mutated *AtHMA4_P_* Δ754Δi harboring MHE1a and MHE1b, we observed no significant change in *GUS* transcript levels or GUS activity (Fig. 4G, H, L, and M, compare >MHE1 with Δ754Δi). The additional introduction of point mutations to generate both copies of MHE2 of *AhHMA4-1_P_* in *AtHMA4_P_* Δ754Δi led to a 4-fold increase in relative *GUS* transcript levels (Fig. 3C; Fig. 4F-H, J, and L, compare >MHE1&2 with >MHE1 and Δ754Δi). A similar effect was obtained by introducing the two copies of MHE2 alone into *AtHMA4_P_* Δ754Δi (Fig. 4F, G, I, J, and L, compare >MHE2 with >MHE1&2 and Δ754Δi). GUS enzyme activity, however, remained low unless the DR was fully deleted (Fig. 4B-M, compare MHE2 > Δ597Δi and > Δ597Δi with >MHE2 and Δ754Δi), consistent with other observations (see Fig. 1F, note the complex effects of the DR of *AtHMA4_P_* on GUS enzyme activity). These data support the enhancing effect of MHE2 on transcript levels in the sequence context of the *A. thaliana* promoter. Other resident *cis*-regulatory functionalities present in *AtHMA4_P_* Δ754Δi may have interfered with the functioning of *A. halleri* MHE1 *cis*-regulatory elements in this sequence context.

**Figure 4.**
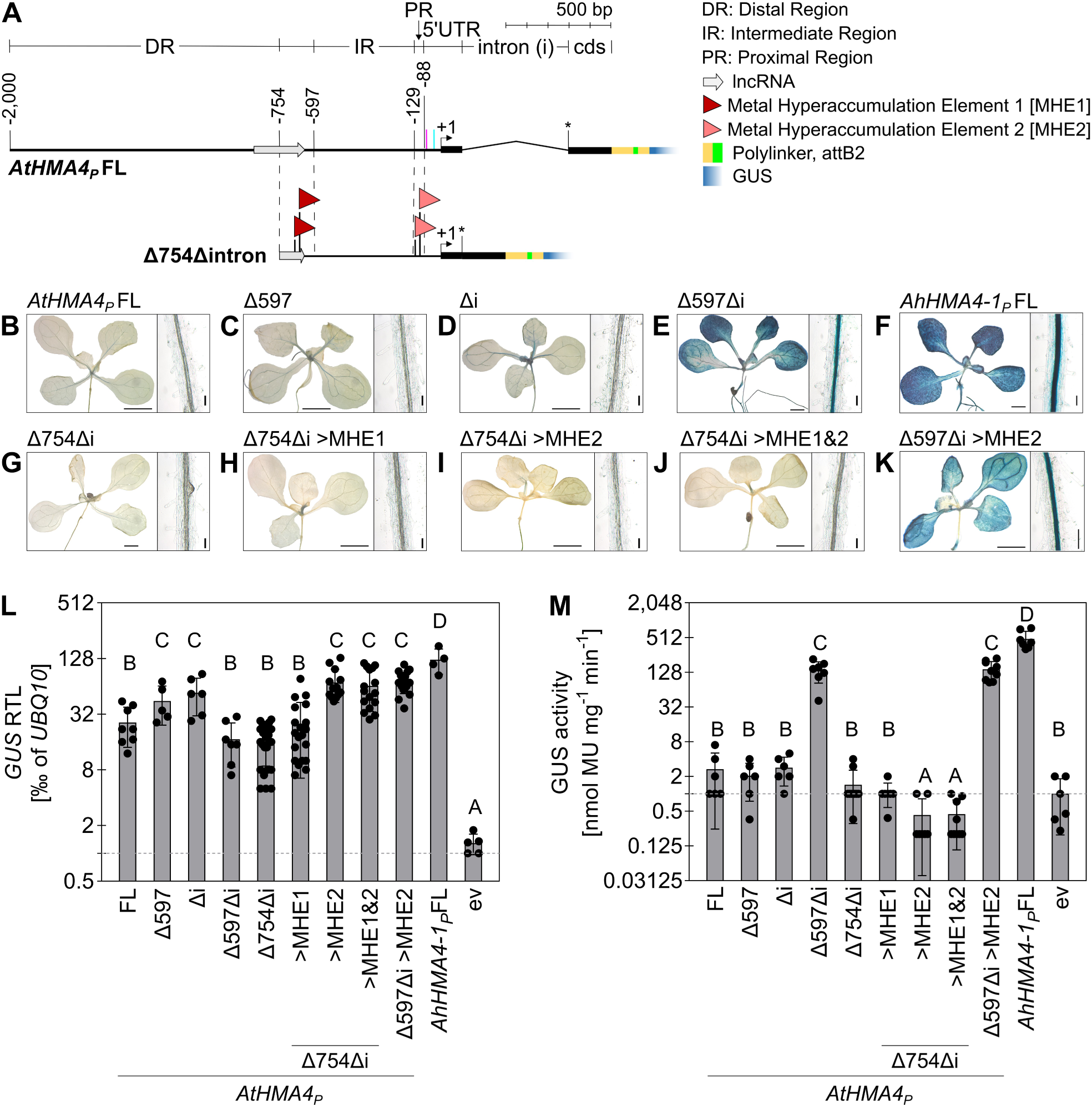
Introduction of two *cis*-regulatory enhancer MHE2 elements is sufficient for conferring enhanced activity to the *AtHMA4* promoter. (A) *AtHMA4_P_* constructs employed for the introduction of *cis*-regulatory elements MHE1 and MHE2 from *AhHMA4-1_P_*. A shortened *AtHMA4_P_* construct (Δ754Δi) that lacks both the intron and 1,246 bp at the 5’ end (corresponding to part of the DR) was used for introducing two copies of MHE1 or of MHE2 by site-directed mutagenesis in positions equivalent to those in *AhHMA4-1_P_* (flag symbols mark the positions). See also Fig. 3B and C. **(B-K)** Histochemical detection of GUS activity in rosettes (left) and the root hair zone of roots (right) in transgenic *A. thaliana GUS* reporter lines. Size bars: 1 mm (E, F, K), 0.5 mm (all other rosette images), 0.2 mm (root images). (L and M) Relative *GUS* transcript levels (L) and specific GUS enzyme activities (M) in seedlings of transgenic *A. thaliana GUS* reporter lines. Bars show mean ± SD (*n* = 3 to 24) of independent transgenic lines. Each datapoint represents the mean of three technical replicates (see Fig. 1). Different characters indicate statistically significant differences based on one-way non-parametric ANOVA followed by a Dunn’s multiple comparison test (*P* < 0.05). >, introduction of MHE1 or MHE2, or of both (MHE1&2) into the Δ754Δi or another construct, as specified. See also Fig. 1B-E.

### *AhHMA4-1* promoter-dependent transcript levels require *CCA1*

We compared MHE1 and MHE2 against databases containing plant transcription factor binding sites using the automated Motif Comparison Tool Tomtom within the MEME suite (Gupta et al. 2007; Bailey et al. 2009). Accordingly, proteins containing MYB and MYB-related domains are the most likely direct interactors of both the MHE1 and the MHE2 *cis*-regulatory elements (InterPro IPR006447; Suppl. Figs. S5B-D and S6B-D, Suppl. Datasets S3 and S4). MHE1 retrieved MYB superfamily proteins of the R2R3 family, comprising two MYB repeats, and of the 3R-MYB family harboring three MYB repeats, as well as transcription factors of the trihelix family that share a domain with some similarity to a single MYB repeat (Kaplan-Levy et al. 2012) (Suppl. Fig. S5C, Suppl. Dataset S3). MHE2 retrieved MYB-related R1R2-subgroup transcription factors that harbor a single R1/R2-type of MYB repeat (Dubos et al. 2010) (Suppl. Fig. S6C, Suppl. Dataset S4).

Among the specific transcription factor proteins identified here, the top-scoring transcription factor proteins predicted to bind MHE2 were in the REVEILLE subamily of circadian clock regulator proteins (Rawat et al. 2009). Target genes of these transcription factors typically show circadian and often also diel oscillations of transcript levels. To test whether *AhHMA4* promoter activities change in a time-of-day-dependent manner, we quantified relative *GUS* transcript levels in our reporter lines over a diel cycle. Relative *GUS* transcript levels driven by *AhHMA4*-*1_P_* to *-3_P_* reached a diel maximum around Zeitgeber Time 9 (ZT9), i.e. 9 h after subjective dawn in the subjective afternoon, at about 1.7- to 2-fold the minimum transcript levels at ZT1 or ZT5, in the *A. thaliana* genetic background (Fig. 5A, and Suppl. Fig. S7A). Transcript levels of *CIRCADIAN CLOCK-ASSOCIATED 1* (*CCA1*), which encodes a central component of the circadian clock in the REVEILLE subfamily of R1/R2 MYB transcription factors, were maximal in seedlings at approximately at ZT1 (Fig. 5B). By contrast, *GUS* transcript levels were much lower and constant over the day in *AtHMA4_P_* reporter lines, and also in lines in which both copies of MHE2 of *AhHMA4-1_P_* were mutated to the corresponding sequence *AtHMA4_P_* (Fig. 5A and Suppl. Fig. S7A, compare *AhHMA4*-*1_P_* to *-3_P_* with MHE2_mut_ and *AtHMA4_P_*, see Fig. 3). These results indicated that *AhHMA4*-*1_P_* to −*3_P_* confer not only strongly elevated transcript levels but also their diel dynamics, both of which depend on MHE2 at least for *AhHMA4-1_P_*, in contrast to *AtHMA4_P_* that lacks MHE2.

**Figure 5.**
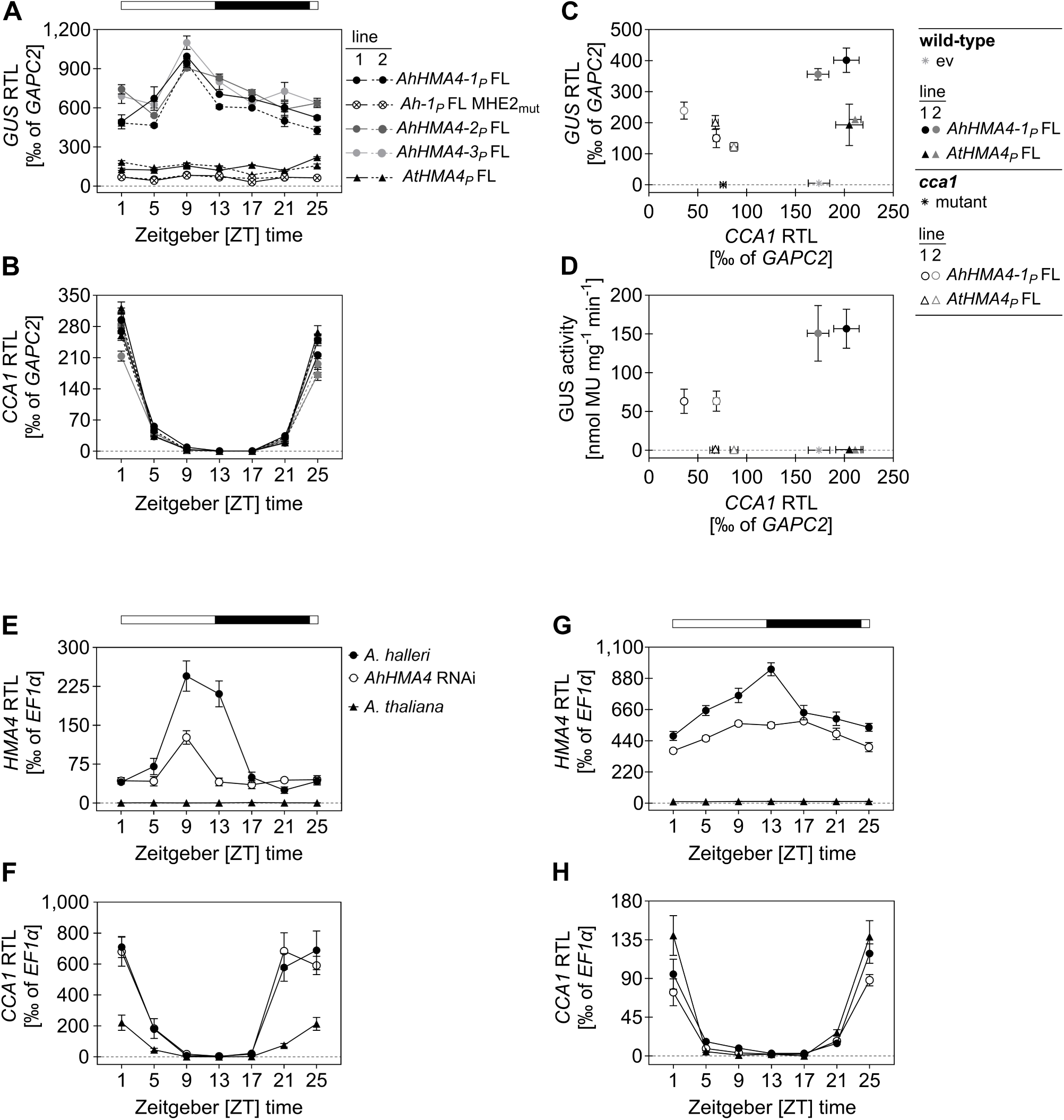
Elevated levels and diel dynamics of *AhHMA4-1_P_*-mediated *GUS* reporter transcript levels depend on both MHE2 and *CCA1* in *A. thaliana*, and diel dynamics of *HMA4* transcript levels in *A. halleri*. (A-D) Relative *GUS* (A) and *CCA1* (B) transcript levels, measured through a diurnal cycle, in seedlings of transgenic *A. thaliana GUS* reporter lines for various *HMA4* promoters. Relative *GUS* transcript levels (C) and specific GUS enzyme activities (D) shown plotted against *CCA1* transcript levels for full-length *AhHMA4-1_P_* and *AtHMA4_P_ GUS* reporter constructs introduced into either wild-type *A. thaliana* plants or into a *cca1-1* loss-of-function mutant (Col-0). Two independent transgenic lines (1, 2) are shown per construct (A-D). Plants were cultivated on agar plates and harvested on day 21 at ZT1 (see also Suppl. Fig. 7). **(E-H)** Diel time course of relative *HMA4* and *CCA1* transcript levels in shoots (E and F) and roots (G and H) of *A. halleri* wild-type plants and one HMA4 RNAi line, as well as in *A. thaliana*. Plants were cultivated in hydroponics and harvested on day 21 (E-H, see also Suppl. Fig. 8). Datapoints are mean ± SD (*n* = 3) of technical replicates from one representative experiment of a total of two independent experiments. Horizontal bar: white fill, day; black fill, night (A, B, E-H). Datapoints (A-H) are mean ± SD (*n* = 3) of technical replicates from one representative experiment of a total of two independent experiments.

Of the transcription factors predicted *in silico* to bind to MHE2 (see Suppl. Fig. S6B to D, Suppl. Dataset S4), we previously observed that *CCA1* transcript levels were 20-fold higher in *A. halleri* than in *A. thaliana* (Becher et al. 2004). We therefore tested whether *CCA1* is necessary for elevated promoter strength of *AhHMA4* promoters and the observed diel oscillations in their activity. We crossed two independent reporter lines of each full-length *AhHMA4*-*1_P_* and *AtHMA4_P_* with a *cca1* mutant line and with a *CCA1* overexpressor line, both of which were previously characterized (Wang and Tobin 1998; Hall et al. 2003) (note that other available and previously characterized lines are not in the Col-0 genetic background). Compared to *AhHMA4*-*1_P_* reporter lines in the wild-type genetic background, *GUS* transcript levels and enzyme activity were reduced to about 55% and 60%, respectively, in *cca1* (Fig. 5C and D). Diel time courses based on RNA extracted from shoots (Suppl. Fig. S7B and C) and roots (Suppl. Fig. S7D and E) further supported a strong *CCA1*-dependence of both the elevated magnitude and the diel changes of transcript levels. By contrast, there was no *CCA1*-dependence of *GUS* expression in *AtHMA4_P_* reporter lines (Fig. 5C and D, Suppl. Fig. S7B-E). Diel dynamics in transcript levels of *GUS* under the control of *AhHMA4*-*1_P_* were also eliminated in a *CCA1* overexpressor background, with little or no change in magnitude (Suppl. Fig. S7F and G). Together, these results suggested that the *cis*-regulatory element MHE2 and CCA1, a transcription factor predicted *in silico* to bind to MHE2, are necessary for both the enhancement and the diel regulation of *AhHMA4-1_P_* promoter activity in the *A. thaliana* genetic background. We also noted that constitutively increased *CCA1* expression alone is not sufficient for further enhancing *AhHMA4-1_P_* promoter activity, which may suggest that the activating role of CCA1 additionally requires one or several other proteins.

### Strong diel dependence of *HMA4* transcript levels in *A. halleri*

Based on our observations in transgenic *A. thaliana* reorter lines, we would expect that *HMA4* trancript levels undergo diel cycles in *A. halleri*. A diurnal time course of *HMA4* transcript levels in shoots and roots confirmed this expectation (Fig. 5E and G, Suppl. Fig. S8A and C). We observed maxima of *HMA4* transcript levels at ZT9 and ZT13 in shoots and roots of *A. halleri*, respectively. As expected, *AtHMA4* transcript levels were low and constant in *A. thaliana*, with *AhHMA4* transcript levels at least 61-fold and up to 1,200-fold higher in roots and at least 43-fold and up to 77-fold higher in shoots of *A. halleri*, depending on the time of day. In shoots, maximum *CCA1* transcript levels were *ca*. 3-fold higher in *A*. *halleri* than in *A. thaliana*, in qualitative agreement with the microarray hybridization-based observations of Becher et al. (2004) (Fig. 5F and Suppl. Fig. S8B). Overall, *CCA1* transcript levels were unaffected in an *A. halleri HMA4* RNAi line, in which *HMA4* transcript levels were strongly suppressed in both shoots and roots compared to the wild type, at least around ZT9 to ZT13, in agreement with Hanikenne et al. (2008). These data demonstrated that *HMA4* transcript levels are under diel regulation in *A*. *halleri* but not in *A. thaliana*, consistent with our promoter analyses in the *A. thaliana* genetic background. Given our finding that high *AhHMA4-1_P_* activity is *CCA1*-dependent in *A. thaliana*, elevated endogenous expression levels of *AhCCA1* in shoots at in the subjective late night and morning (ZT21, 25, 1 and 5) may contribute to enhanced transcript levels of *AhHMA4* and their diel dynamics in *A. halleri*.

## Discussion

### The *cis*-regulatory changes in *HMA4* during the evolution of metal hyperaccumulation in *A. halleri*

Compared to *AtHMA4*, the promoter activities of the three tandem *AhHMA4* gene copies present in *A. halleri* are enhanced, thus resulting in strongly elevated transcript levels of *AhHMA4*, the key locus making the largest contribution to metal hyperaccumulation and hypertolerance in *A. halleri* (Talke et al. 2006; Courbot et al. 2007; Hanikenne et al. 2008). Here we identify the causal c*is*-regulatory sequence differences contributing to *AhHMA4* promoter activity, namely the enhancer elements MHE1 and MHE2 which are predicted to bind R2R3 MYB and R1R2 MYB-related transcription factors, respectively (Figs. 1-4, Suppl. Figs. S1 to S6). Deleting or mutating the MHE1 and MHE2 elements in *AhHMA4-1_P_* either alone or in combination was consistent with their additive effects on the transcription of the downstream gene (Figs. 1 and 3). With no exceptions, the MHE1a, MHE2a and MHE2b elements were present in microsyntenic promoter regions of the three *HMA4* gene copies of all *A. halleri* accessions for which data are presently available (Hanikenne et al. 2013) (Suppl. Figs. S5 and S6). Upon introduction of these *cis*-regulatory elements into an *AtHMA4* promoter context by site-directed mutagenesis, our results suggested that MHE2 is sufficient for enhancing transcript levels (Fig. 4). It is possible that we were unable to detect the enhancing effect of MHE1 introduced into *AtHMA4_P_* because of a specific repressive context therein. We detected repressive functions in the DR of the *AtHMA4* promoter, and the combination of both MHE1 and MHE2 within *AtHMA4_P_* did not confer as high levels of reporter gene transcripts as did full-length *AhHMA4-1_P_* (Fig. 4). Alternatively, it could be that MHE1 has no functional relevance for the differences in promoter activity between the *A. halleri* and *A. thaliana HMA4* promoters despite functioning as an enhancing *cis*-regulatory element in *AhHMA4-1_P_*. Support for this alternative hypothesis may be drawn from the fact that when both MHE1 and MHE2 were mutated in *AhHMA4-1_P_*, reporter gene transcript levels were well below those observed in *AtHMA4_P_* lines (Fig. 3).

Our results suggested that MHE2 in the PR of the *AhHMA4-1_P_* region is both necessary and sufficient for elevated levels of promoter activity above those of unmodified *AtHMA4_P_* (Figs. 3-5). Thus, we conclude that MHE2 binds a transcription factor which mediates the activation of transcription of the downstream gene either directly or indirectly. In all individuals of *A. halleri*, we identified two copies of MHE2 (8 bp) in the PR of each of the three *HMA4* gene copies, spaced by 17 (*AhHMA4-1_P_*), 18 (*AhHMA4-3_P_*), or 27 to 30 bp (*AhHMA4-2_P_*) (Suppl. Fig. S6). The conservation of both MHE2 elements and identical diel rhythmicity of *AhHMA4-2_P_*- and *AhHMA4-3_P_*-directed promoter activity suggested that these findings are valid for the promoters of all three *HMA4* gene copies in *A. halleri* (Fig. 5, Suppl. Figs S7 and S8). There was some degree of sequence variation between the two copies of MHE2 in the PR of each *HMA4* gene and among the three *HMA4* gene copies of each genotype. By contrast, there was hardly any polymorphism between *A. halleri* individuals originating from Europe, consistent with the earlier analysis of broader promoter regions (Hanikenne et al. 2013). Three out of four alleles of an *AhHMA4-3*-like promoter sequence of *A. halleri* from the Tada mine (Japan) carried point mutations in MHE2b, which could be related to local selection pressure for nutrient balancing and deserves further study (Hanikenne et al. 2013; Stein et al. 2017; Krämer 2024).

### MHE2 directs both elevated and rhythmic *AhHMA4* promoter activity in dependence on *CCA1*

The consensus MHE2 motif (CACTATCT) was reminiscent of the evening element (EE, consensus AAATATCT). EEs confer circadian clock-regulated gene expression and are well-known as target sites for CCA1 binding (Wang et al. 1997; Nagel et al. 2015; Kamioka et al. 2016). EEs can also interact with other clock oscillator proteins such as LHY and RVE8, all of which are members of the same REVEILLE family of MYB-related R1/R2-type proteins in *A. thaliana* (Rawat et al. 2009). In the *AhHMA4-1_P_* sequences of all European *A. halleri* accesions, MHE2a was identical to the consensus EE, whereas the sequence of MHE2b was TAATATCA.

In a *cca1* mutant, *AhHMA4-1_P_*-driven reporter transcript levels were no higher than *AtHMA4_P_*-driven reporter transcript levels (Fig. 5), and their diel rhythmicity was lost (Suppl. Fig. 7), very similar to the effects of mutating the MHE2 elements in *AhHMA4-1_P_*. This supported the *CCA1*-dependence of the difference between *A. halleri* and *A. thaliana HMA4* promoter strengths and provided circumstantial evidence for MHE2 as a target *cis*-regulatory element for CCA1-dependent transcriptional activation. Moreover, the similarity between MHE2 and the EE is consistent with a model of direct binding of CCA1 to MHE2 for causing or participating in the activation of transcription of the downstream reporter gene or *AhHMA4*.

Earlier microarray-based cross-species transcriptomics identified *CCA1* as a constitutively more highly expressed gene in shoots of *A. halleri* by comparison to *A. thaliana* (Becher et al. 2004). These findings are confirmed here (Fig. 5F), thus providing some additional circumstantial support for the proposed regulatory role of *CCA1* in the expression of *HMA4* in *A. halleri*. The literature addresses CCA1 as both an activator and a repressor of transcription, with reports of the latter function being more widespread and known particularly for the regulation of genes encoding components of the evening loop in the core circadian oscillator of *A. thaliana* (Wang et al. 1997; Wang and Tobin 1998; Alabadí et al. 2001; Farré et al. 2005; Lu et al. 2009; Dong et al. 2011; Dong et al. 2011; Nagel et al. 2015; Kamioka et al. 2016). Consequently, CCA1, which contains no characterized activation or repression domains, is considered to possess alternative activating or repressing functionalities dependent on the regulatory context including its interacting proteins, but the mechanistic basis of this remains poorly understood (Nagel et al. 2015; Kamioka et al. 2016). CCA1 and LHY often act as heterodimers (Lu et al. 2009), but these are apparently not required for normal clock functioning (Kamioka et al. 2016).

The *cca1* mutant has only mild phenotypes, including a shortened circadian period with respect to the transcript levels of core circadian oscillator genes, as well as early flowering (Green and Tobin 1999; Mizoguchi et al. 2002). By comparison, the strongly lowered *AhHMA4-1_P_*-driven reporter gene expression in the *cca1* mutant observed in this study (Fig. 5C) supports a role for *CCA1*, but not for *LHY* at this point, in *AhHMA4-1_P_*-regulated reporter gene transcript levels.

In this study, *CCA1* transcript levels peaked around ZT1, whereas *AhHMA4_P_*-directed reporter gene transcript levels in *A. thaliana* and *HMA4* transcript levels in *A. halleri* peaked considerably later, at ZT9, in shoots (Fig. 5E and 5F). Transcript levels of previously proposed direct target genes of CCA1-dependent transcriptional activation were reported to peak in the late morning to afternoon under diel light-dark cycling conditions (Wang and Tobin 1998; Farré et al. 2005). Accordingly, transcript levels of *LHCB1.1*/*CAB2* are maximal around ZT4 and remain high until ZT8, although CCA1 protein was detectable only between ZT22 and ZT4 and no longer at ZT8 (Wang and Tobin 1998). Transcript levels of direct CCA1 targets *PSEUDO-RESPONSE REGULATOR 9* and *7* encoding circadian clock transcription factors peaked around ZT4 (PRR9) and ZT8 (PRR7) under diel cycling conditions (Farré et al. 2005). CCA1 binds to the *PRR5* promoter, and whereas CCA1-FLAG protein levels were detectable between ZT0 and ZT6, *PRR5* transcript levels peak around ZT9 under diel cycling conditions (Kamioka et al. 2016). The authors noted that this delay may reflect the required time for the *PRR5* transcript to accumulate, or alternatively suggest that CCA1 represses *PRR5* transcription in the morning. Since *PRR5*, *PRR7* and *PRR9* form part of the complex interacting regulatory loops of the circadian clock, changes in their transcript levels in circadian clock mutants are generally difficult to interpret. Transient assay systems suggested that CCA1 alone has at least an immediate repressive effect on their transcript levels (Kamioka et al. 2016). Finally, CCA1-mediated indirect regulation of target genes could occur *via* effects on chromatin structure, as was proposed for the repressive effects of CCA1 (Perales and Más 2007).

### Comparison with *HMA4* promoters of other Zn/Cd hyperaccumulator and non-accumulator Brassicaceae species

Our results suggest that two EE-related *cis*-regulatory elements (MHE2) present in the promoter of each of the three *A. halleri HMA4* gene copies confer their regulation through the pre-existing circadian clock transcriptional network, different from *A. thaliana HMA4* that lacks MHE2 elements. A single MHE2 copy resembling the EE is present in the PR of the *HMA4* promoter of the non-hyperaccumulator species *A*. *lyrata*, of which leaves contain 2-fold higher *HMA4* transcript levels compared to *A. thaliana* (Hanikenne et al. 2013) (Suppl. Fig. S6). Similar to the presence of MHE2 elements in the PR of *AhHMA4* promoters, we also identified *in silico* predicted MHE2 elements in the promoters of the four *HMA4* gene copies in the Zn/Cd hyperaccumulator *Noccaea caerulescens*, *NcHMA4-1* to *4-4* (Suppl. Datasets S5 and S6). *N. caerulescens* is in the phylogenetically distant lineage II of the Brassicaceae family that diverged from the *Arabidopsis* species in lineage I more than 20 Mya (Clauss and Koch 2006). Thus, in addition to convergent *HMA4* gene copy number expansion, convergent promoter mutations may have occurred in this species to result in elevated *HMA4* transcript levels (Lochlainn et al. 2011). The *HMA4* promoter region of *A. lyrata* also harbors an MHE1 element in a microsyntenic position (Suppl. Fig. S5, Suppl. Dataset S5). We additionally observed MHE1 elements in the distal promoter regions of all four *HMA4* gene copies of *N. caerulescens* (Suppl. Fig. S5) (Lochlainn et al. 2011). By contrast, all *A. thaliana* accessions and *Capsella rubella* in lineage I, as well as *Brassica rapa* and *Arabis alpina* in lineage II, lack a precisely identical copy of MHE1, or MHE2 copies in their proximal promoter regions entirely (Suppl. Datasets S5 and S6).

According to our reporter lines, complex repressive functionalities in the DR of *AtHMA4_P_* and in the intron in the 5’ untranslated region of *AtHMA4* appear to act not only at the transcript level, but possibly additionally at the protein level (Fig. 1, Suppl. Fig. S3). The DR contains an annotated lncRNA (−859 to −621; Suppl. Fig. S3). The sequence features and molecular mechanisms involved in the *cis*-repression of *AtHMA4* will require further study in the future. Our analysis did not detect any evidence for these repressive functionalities acting on any *A. halleri HMA4* gene copies. Either *A. halleri* lost these repressive functionalities through mutations, or *A. thaliana* acquired these repressive functionallities only recently, after the split of the *A. thaliana* and *A. halleri* lineages. All in all, our data suggest that the former is more likely than the latter (Suppl. Fig. S3, Suppl. Datasets S5 and S6). Irrespective of this, we cannot exclude the possibility that some degree of *cis*-activation of *HMA4* may have been the ancestral state in lineages I and II of the Brassicaceae.

### Co-option of gene regulatory networks through *cis*-regulatory mutations in *HMA4*

Gene regulatory network co-option is often exclusively conceptualized as involving *trans* regulatory change, following the rationale that one such mutation can alter the expression of multiple target genes downstream in the regulatory hierarchy (True and Carroll 2002; Eden McQueen and Rebeiz 2020). By comparison, *cis*-regulatory mutations at multiple loci in the genome would be required to achieve an equivalent alteration in overall gene expression, which is far less likely to occur as a result of evolutionary processes. Yet, there are examples of *cis*-regulatory mutation-based co-option in evolutionary novelties, although the underlying sequence elements have largely remained elusive. Indeed, *cis*-regulatory change could be largely equivalent to changes in *trans* in cases where a *cis*-regulatory mutation adds a new expression domain for an existing transcription factor, for example. Past research and theoretical considerations have almost exclusively considered morphological novelty and developmental contexts, however, whereas this present study addresses physiological novelty. Here, *cis*-regulatory change change results in the co-option of a regulatory network upstream in the hierarchy to enhance product dosage of the *HMA4* gene encoding a pump which exports heavy metal cations, primarily Zn^2+^ and Cd^2+^, from specific cell types. This has the known fitness benefits of affording elemental defence against herbivory and enhanced heavy metal tolerancee and thus fitness on soils containing high, toxic levels of Zn and Cd. It remains to be invistigated whether the diel rhythmicity of *HMA4* transcript abundance is mirrored at the HMA4 protein level. Although membrane transport proteins are no classical *trans* regulators, altered expression of *HMA4* can affect the expression of other genes in *trans*, as was demonstrated in both *A. halleri* and *A. thaliana* (Hanikenne et al. 2008; Sinclair et al. 2018). In *A. halleri HMA4* RNAi lines, for example, transcriptional Zn deficiency responses were suppressed through the Zn homeostasis network as a result of locally increased intracellular Zn levels (Sinclair and Krämer 2012; Krämer 2018). The affected genes are among the candidates thought to contribute to the metal hyperaccumulation syndrome in *A. halleri* wild-type plants, based on their strongly elevated transcript levels compared to *A. thaliana*. Thus, the *cis*-regulatory change of *AhHMA4* acts homeostatically in *trans* to additionally co-opt the Zn homeostasis gene regulatory network. While the co-option of components within the metal homeostasis network might restrict the flexibility of the system, several factors — such as the homeostatic nature, and generally network redundancy, modularity, and the partial nature of co-option — can reduce the likelihood of detrimental effects on metal homeostasis (Eden McQueen and Rebeiz 2020). Notably, *HMA4* transcript levels are not Zn-deficiency responsive in either *A. halleri* or *A. thaliana* (Talke et al. 2006). Thus, the Zn homeostasis network can maintain its core functions while still allowing for evolutionary innovation through its co-option. Homeostatically generated gene expression changes governed by *HMA4* gene product dosage allow for continued plasticity in the extreme physiological traits of metal hyperaccumulation and associated hypertolerance, possibly differing from the co-option of transcription factors implementing specific developmental modules.

### Conclusions

In conclusion, our results suggest that *cis*-regulatory mutations generating two tandem copies of MHE2 were necessary and sufficient for enhanced activity of the *AhHMA4-1* promoter. The same *cis*-regulatory elements are likely to be effective in the other two *HMA4* gene copies as a result of gene duplication events in *A. halleri*. *CCA1* is neccesary for both elevated levels and diel cycling of *AhHMA4-1_P_*-regulated transcript levels. By contrast, there is no diel or *CCA1*-dependent regulation of the generally low *AtHMA4* promoter activity. Instead, our results suggest that *AtHMA4* expression undergoes complex repressive regulation, thus contributing to the *cis*-regulatory divergence between *HMA4* genes of *A. halleri* and *A. thaliana*.

## Methods

### Plant material and growth conditions

Seeds of the *A. thaliana* wild type (Col-0) were from were obtained from Lehle seeds (Round Rock, TX, USA). Seeds of *A. thaliana cca1-1* (N67781, *cca1-1* mutant N67780 in the Ws background was backcrossed six times with Col-0 to generate this stock) carrying a T-DNA insertion in the fourth intron of *CCA1* (Krysan et al. 1996; Green and Tobin 1999; Yakir et al. 2009) and of a transgenic line expressing *CCA1* (*CCA1*-OX o38, N67794 in the Col-0 background) under the control of the Cauliflower Mosaic Virus 35S Promoter (Wang and Tobin 1998) were purchased from the Nottingham Arabidopsis Stock Centre (NASC). After obtaining them from NASC, homozygous lines were confirmed through late flowering (*CCA1*-OX o38) (Wang and Tobin 1998), and PCR-based genotyping following DNA extraction (Edwards et al. 1991) (*cca1-1*) (Suppl. Table S1 for details about primer pairs used for the genotyping of the *cca1-1* mutant lines). Transgenic *A. thaliana GUS* reporter lines for full-length *HMA4* promoters of *A. halleri* and *A*. *thaliana* (L. Heynhold, accession Col-0) were from Hanikenne et al. (2009) and Nouet et al. (2015) (*HMA4-2* and *4-3*). The *AtHMA4_P_* construct comprised a genomic fragment of 2,799 bp in length that included the initial 204 bp of *AtHMA4* coding sequence, corresponding to 68 amino acids (aa) predicted to be cytosolic, with the first out of a total of 8 conserved transmembrane helices predicted to include amino acids 95 to 115; TmConsens, https://aramemnon.botanik.uni-koeln.de/). *Arabidopsis halleri* (L. O’Kane and Al-Shehbaz) ssp. *halleri*, population Langelsheim, accesssion Lan3.1, was collected in the field and maintained in the greenhouse on soil (Minitray soil, Balster Einheitserdewerk, Frӧndenberg) through vegetative propagation (Becher et al. 2004). A previously charcaterized *AhHMA4* RNA interference (RNAi) line (4.2.1) was used here (Hanikenne et al. 2008).

For propagation of homozygous lines and/or genotyping, *A. thaliana* were grown on soil after stratification (at 4°C for 3 days) in square pots (5 x 5 x 5 cm) in a glasshouse (16-h-day at 20 to 22°C, with daylight supplemented by sodium vapour lamps to an intensity of 120-150 µmol m^-2^ s^-1^ when necessary, night 18 to 20°C). For sterile cultivation, seeds were sterilized in 70% (v/v) ethanol for 5 min, followed by 10 min in 0.65 to 0.78% (w/v) NaOCl containing 0.05% (v/v) Tween 20 in ultrapure water, followed by 4 washes in sterile ultrapure water. Initial screening in the T1 generation was done by sowing 50 mg sterilized seeds (approximately 2,500 seeds) per line on Type M agar-solidified (0.8% w/v) 0.5x MS medium supplemented with 30 µg ml^-1^ of Hygromycin B (Hyg) from Duchefa (catalog no. H0192, Haarlem, NL) alongside a negative (Col-0) and a positive T3 homozygous control.

For the characterization of promoter-reporter lines, sterilized seeds were positioned along a horizontal line at 2.0 cm distance from the upper edge across a 120 mm × 120 mm square polystyrene Petri plate (Greiner Bio-One GmbH, Frickenhausen, Germany) onto 50 ml of an agar-solidified (0.8% (w/v), Type M, Sigma-Aldrich, Steinheim, Germany) modified Hoagland solution (see below) supplemented with 1% (w/v) sucrose. Plates were stratified in darkness at 4°C for 48 to 72 h, followed by transfer into growth chambers (CFL Plant Climatics, Wertingen, Germany) for cultivation in vertical orientation (with the line positioned horizontally across the upper end) in a 12-h day (120 μmol m^−2^ s^−1^, 22°C) / 12-h night (18°C) regime. For RNA and protein isolation, whole seedlings, or root and shoot tissues separated with a scalpel, were harvested at ZT1 on day 21, tissues were pooled from four replicate plates (80 plant individuals in total) per line, frozen in liquid nitrogen and stored in 50-mL screw-cap polypropylene tubes at −80°C until further processing. For diel time course experiments of *A. thaliana* lines seedlings were harvested at ZT1 on day 21 and until of ZT25 of day 22, and processed as described above.

For diel time course experiments including *Arabidopsis halleri*, we used accession Lan3.1 as the wild type, and *AhHMA4* RNAi line 4.2.1 (Hanikenne et al. 2008). Vegetative cuttings of a minimum of 7 cm in length were prepared from mother plants cultivated on soil (Minitray soil mixed with 5% (v/v) soil from Langelsheim field site; Stein et al. 2017) in 2-l pots, and transferred into a 50-ml polypropylene tube containing modified Hoagland solution (see below) for rooting and pre-cultivation. *A. thaliana* Col-0 seedlings were pre-cultivated in sterile conditons as described above. For hydrophonic cultivation, 21-d-old *A*. *thaliana* seedlings and 17-d-old *Arabidopsis halleri* clones were transfered into 400-ml vessels containing modified Hoagland solution (0.28 mM KH_2_PO_4_, 1.25 mM KNO_3_, 1.5 mM Ca(NO_3_)_2_, 0.75 mM MgSO_4_, 5 µM of a complex of Fe(III) and N,N′-di-(2-hydroxybenzoyl)-ethylenediamine-N,N′-diacetate (HBED), 25 µM H_3_BO_3_, 5 µM MnSO_4_, 5 µM ZnSO_4_, 0.5 µM CuSO_4_, 50 µM KCl, and 0.1 µM Na_2_MoO_4_, buffered to pH 5.7 with 3 mM 2-(N-morpholino)ethanesulfonate) in ultrapure water. Roots and shoots were harvested every 4 h through a 24-h cycle beginning at ZT1 on day 21 and until ZT25 of day 22 of cultivation.

### DNA cloning

Promoter deletion constructs were made in groups according to the procedures described next (designated 1 to 3; Suppl. Table S1, Suppl. Table S2). (1) The first set of promoter deletion constructs were initially generated in the pBluescript II KS+ (pBKS) vector backbone (X52327.1, NCBI) for the transient transfection of *A. thaliana* protoplasts. Prior to this, the GW-GUS-tNOS cassette of the pMDC163 destination vector (4,124 bp) (Curtis and Grossniklaus 2003), which contains a gateway (GW) recombination cassette, the *uidA* (*GUS*) reporter gene, and a nopaline synthase terminator (tNOS), was amplified by PCR using primers containing added 5’ *Xho*I restriction sites, followed by its cloning into the corresponding site in the pBKS plasmid. Second, full-length *AtHMA4_P_* or *AhHMA4*-*1_P_* to *AhHMA4*-*3_P_* fragments (Hanikenne et al. 2008) were recombined from pENTR-D/TOPO entry clones into the pBKS-GW-GUS-tNOS vector using the Gateway® LR Clonase™ II enzyme mix kit from Invitrogen. Third, the deletions were introduced in the full-length promoter constructs using PCR-based site-directed mutagenesis (Wang and Malcolm 1999). Deletion primers were designed to flank the 5’ and/or 3’ extremities of the deleted segments, respectively (Suppl. Table S2) and used for linear PCR amplification of the pBKS-*HMA4_P_*-GW-GUS-tNOS vectors (14 to 18 cycles of 95 °C for 30 sec, 55 °C for 60 sec, 68 °C for 16 min), followed by digestion of the parent DNA with *Dpn*I, and subsequent transformation of *E. coli*. All PCR reactions were performed using a proofreading polymerase (Pfu Turbo, from Stratagene), and verified by Sanger sequencing. In GUS assays following transient transfection with the pBKS constructs in various systems, we observed that GUS enzyme activities were poorly reproducible across technical replicates and independent experiments. Consequently, all constructs were then subcloned into the binary plasmid pMDC163 (Curtis and Grossniklaus 2003) for the stable transformation in *A. thaliana* (see 2 below). (2) Another set of deletion constructs was made within the pBKS-*HMA4_P_*-GW-GUS-tNOS plasmid (Suppl. Table S2). Plasmids generated according to (1) above in the pBKS backbone were either linearized by an *Asc*I/*Pac*I double digestion or by PCR amplification (Herculase II Fusion DNA Polymerase, Agilent). In parallel, appropriate primers were used to PCR-amplify promoter fragments and produce overlapping ends with the pBKS backbone (Herculase II; Suppl. Table S2). Following gel purification, plasmid and promoter fragments were fused (In-Fusion, Clontech) following the manufacturer’s instructions. Finally, the *HMA4_P_*-GW-GUS cassettes were excised from the pBKS vector using *Pac*I/*Sac*I digestion and cloned into the corresponding sites of the pMDC163 vector, generating the final constructs. (3) Promoter-deletion and mutated versions were generated by either (a) standard PCR-mediated amplification, (b) overlap extension PCR (Heckman and Pease 2007), or (c) quick change site-directed mutagenesis (Zheng 2004), employing non-linearized plasmids containing full-length *AtHMA4_P_* or *AhHMA4*-*1_P_*, *AhHMA4-2_P_* or *AhHMA4*-*3_P_* fragments (Hanikenne et al. 2008) using the Q5^R^ high fidelity DNA polymerase from New England BioLabs_INC_ (NEB, product M0515S, Ipswich, Massachusetts, United States) (Suppl. Table S2). For (c), in which GATEWAY destination vectors pMDC163 were used to introduce the desired mutatios, resulting constructs were introduced into *Agrobacterium tumefaciens* GV3101 (see below). For (a) and (b) the resulting promoter fragments were cloned into pENTR/D TOPO (Invitrogen^TM^ product K240020), and One Shot™ TOP10 Chemically Competent *E. coli* cells (ThermoFisher Scientific) were transformed by heat-shock (42°C for 30 s) and selected on LB medium containing 50 µg ml^-1^ kanamycin following manufacturer’s instructions. Plasmid DNA prepared from these entry clones was used for site-directed recombination into the GATEWAY destination vector pMDC163 (Curtis and Grossniklaus 2003), which contains the *uidA* (*GUS*) reporter gene using the the Gateway® LR Clonase™ II enzyme mix kit from Invitrogen (Catalog number: 11791020, Carlsbad, CA USA) following manufacturer’s instructions, with the following modifications. After terminating the LR reactions by the addition of proteinase K, 2.5 units of the *Hpa*I restriction enzyme (NEB, Catalog no. R0105S) were pipetted in each of the LR reactions, following by incubation at 37°C for 3 h. This step was performed to eliminate unrecombined and contaminating pENTR-D/TOPO vector that contains the same bacterial resistance marker as the pMDC163 vector (i.e. *Kan*). Inactivation of the *Hpa*I enzyme was done at 65°C for 20 min. Of these reactions, 1 µl was used to transform One Shot™ TOP10 *E. coli* cells as described above. For entry and destination plasmids, screening of up to 20 colonies was done by colony PCR (Bergkessel and Guthrie 2013) in a total volume of 20 µl using the ThermoScientific DreamTaq Green PCR Master Mix (2x) per colony in combination with 1 µl primer mix of 10 µM forward M13 (5’-GTAAAACGACGGCCAG-3’) and reverse M13 (5’-CAGGAAACAGCTATGAC-3’) using the following PCR program: 95°C for 10 min, 35 cycles of 95°C for 30 s, 55°C for 30 s, 72°C for 1 min per kb, and a final extension at 72°C for 10 min. For colonies yielding PCR products of the correct size, we verified plasmid DNA restriction patterns and finally conducted Sanger sequencing using the forward and reverse M13 universal primers for each entry and each final destination plasmids. pMDC163 plasmids carrying the correct promoter fragments were introduced into *Agrobacterium tumefaciens* GV3101 by electroporation using a MicroPulser electroporator from Bio-Rad (Xu and Qingshun 2008).

### Plant transformation and selection of homozygous lines

Stable transformation of *A. thaliana* was performed using the floral dip method (Clough and Bent 1998). Per construct, we transferred twenty T1 plants onto soil and collected the T2 seeds of each plant individually. At least 120 to 250 T2 seeds for each line were sterilized, stratified and sown on selective medium, as described above. Segregation ratios were determined by counting Hyg-resistant transgenic and sensitive wild-type seedlings. For each line exhibiting a segregation ratio consistent with T-DNA insertion at a single locus (75% Hyg-resistant seedlings) according to a *X*^2^ test, we transferred twelve tolerant seedlings onto soil and collected seeds from each plant. Segregation ratios and *X*^2^ tests were repeated in subsequent generations (typically T3) until obtaining homozygous lines (100% Hyg resistance).

### Crossing and selection of homozygous lines

*A. thaliana* homozygous *cca1-1* mutant and *CCA1*-OX o38 transgenic plants were used as pollen acceptors for crossing with two independent *GUS* reporter lines for each *AtHMA4_P_* and *AhHMA4-1_P_* (pollen donor), as well as *AhHMA4-2_P_* and *AhHMA4-3_P_* (for *CCA1*-OX o38 only) (Weigel and Glazebrook 2006). Five to ten inflorescences were pollinated, marked with colored threads, and siliques were harvested individually 21 d later. F1 seeds were germinated on 50 µg ml^-1^ Kan, and resistant plants were transferred to 30 µg/mL of Hyg after 14 d alongside wild-type Col-0 plants, of the same age, as controls. Plants resistant to both antibiotics were transferred to soil, and their F2 seeds were harvested, sown, and segregation ratios were quantified separately on either 50 µg ml^-1^ Kan or 30 µg ml^-1^ Hyg, confirming single T-DNA insertions, until obtaining lines homozygous for both markers (see above). In addition to this, the presence of the *cca1-1* mutant allele, and of the *GUS* transgene were verfied by PCR (see Suppl. Table S1).

### RNA extraction, cDNA synthesis, and real-time quantitative PCR

Frozen plant tissues were homogenized in 50-ml polypropylene tubes (Avantor Performance Materials GmbH, Griesheim, Germany, Catalog no. 734-0453) pre-chilled in liquid N_2_, each containing a commercial glass marble, using a vortex (Vortex-Genie 2, Scientific Industries, Inc., Bohemia, New York, USA) at maximal speed for 10 s, with three to four cycles of grinding and chilling in liquid N_2_. RNA was isolated from aliquots of frozen homogenized tissue using the Trizol^TM^ reagent from Invitrogen^TM^ (Catalog no. 15596018, Dreieich, Germany). One hundred mg of plant tissue was transferred into a 2-ml polypropylene reaction vial, and 1 mL of Trizol^TM^ was added to each sample, followed by vortexing at maximal speed for 30 s and incubation at root temperature (RT) for 5 min. Samples were centrifuged in a fixed-angle, pre-cooled (4°C), rotor at 18,500 x*g* for 10 min. The supernatant (1 ml) was transferred to new 1.5-ml reaction vial containing 200 µl chloroform (molecular biology grade), and vortexed at maximum speed for 15 s, followed by centrifugation at 4°C at 18,500 x g for 15 min. This step was repeated once in a fresh 1.5-ml reaction vial containing 200 µl of chloroform. Following centrifugation, 450 µl of the supernatant were transferred into a 1.5-ml reaction vial containing 500 µl 2-Propanol (≥99.5%, for molecular biology, Fisher BioReagents™, Darmstadt, Germany, Catalog no. I9516-1L) pre-cooled on ice. Samples were mixed by inverting 5 times, followed by incubation at RT for 10 min. Samples were then centrifuged at 4°C 18,500 x g for 15 min. The supernatant was carefully decanted, and pellets were washed twice in ice-cooled 1 ml of 75% (v/v) ethanol (99% absolute ethanol, Extra Pure, SLR, Fisher Chemical™, Dreieich, Germany) diluted in milli-Q water pre-treated with diethylpyrocarbonate at a final concentration of 0.1% (v/v) followed by vortexing at maximal speed for 30 s. After the final wash, the supernatant was carefully pipetted off, and the pellet was air-dried under the fume hood on a paper towel for 10 min. The RNA was resuspended in 50 μl of nuclease-free water containing 0.1 mM EDTA (pH = 7.2) based on the manufacturer’s recommendations for TRIzol™, followed by incubation on ice for 30 min. RNAs were stored at −80°C until use.

Prior to cDNA synthesis, quality and quantity of RNA were visually assessed by denaturing gel electrophoresis (Mansour and Pestov 2013) and by photometric analysis (A_260_ and A_280_) (Nanodrop 2000/2000c, ThermoScientific^TM^, Dreieich, Germany).

Accordingly, based on manufacturer’s instructions 10 µg of RNA per sample were used for the digestion of contaminating genomic DNA using 3 units of the DNase I (RNase-free) from NEB (product M0303L) in a 150 µl final volume reaction and incubate at 37°C for 30 minutes. Following this step, samples were treated with phenol-chloroform based RNA purification using the Aqua-Phenol reagent (Carl Roth®, Karlsruhe, Germany, Catalog no. A980.3). Samples were then procesed as described for the Trizol^TM^ protocol above, with air-drying for 5 min and a resuspension in a final volume of 30 μl nuclease-free water. RNA yield was quantified spectrophotometrically using a NanoDrop™ 2000/2000c by measuring three replicate 2-µl aliquots of each sample and calculating the average. cDNA was synthesized from 1 µg total RNA in a total volume of 20 µl using the RevertAid First Strand cDNA Synthesis Kit following supplier’s instructions (Thermo Scientific™, Dreieich, Germany, Catalog no. K1622), primed by oligo(dT)_18_. Prior to the storage of cDNA samples at a −20 °C, aliquots consisting of 1-µl of cDNA and 99-µl of nuclease-free water (pH 7) were prepared and used for RT-qPCR (see below).

### RT-qPCR

Primer design for RT-qPCR was done using the LightCycler Probe Design Software version 2.0 (1.0.R.36) (Roche, Mannheim, Germany) with default options (Suppl. Table S1). Real-time PCR reactions were performed in 384-well plates using the PROMEGA GoTaq® qPCR Master Mix (Walldorf, Germany, Catalog no. A6002) on a LightCycler480 (Roche, Mannheim, Germany) to monitor cDNA amplification as described (Quintana et al. 2022). Unless specified in figure legends, every cDNA sample was run in triplicate on a single plate and three independent replicate plates were run for each sample. Reaction efficiencies (E) for each PCR reaction were automatically determined using the LinRegPCR program, version 2016.0 (Ruijter et al. 2009).

Transcript level (TL) of both the constitutively expressed reference and the target genes were calculated as TL = E^−Ct^, where C_t_ represents the cycle threshold. Per replicate plate, RTL was calculated by dividing mean (of the triplicate reactions run in a single plate) TL of a given target gene by mean (same as for the target gene) TL of the reference gene. RTL of *GUS* driven by various *HMA4* promoter fragments were independently normalized to each of four of the most stable constitutively expressed reference genes in *A. thaliana*, *ACT2*, *EF1α*, *GAPC2*, and *UBQ10* (Czechowski et al. 2005). Reaction efficiencies and independent C_t_ values calculated for each reference gene and the *GUS* reporter gene (Suppl. Dataset S7). The mean PCR reaction efficiency of *UBQ10* were the most consistent across all 429 qPCR reactions, and the average C_t_ values for *UBQ10* were the most consistent compared to the other three housekeeping genes. Furthermore, across 106 independent *A. thaliana* transformants there was a positive correlation of specific GUS enzyme activity to relative *GUS* transcript levels normalized to *UBQ10* (*R*^2^ = 0.74) with the highest correlation coefficients (Suppl. Dataset S7). Consequently, we selected *UBQ10* as the reference gene for the calculation of relative *GUS* transcript levels at single time points in *A. thaliana*. In Suppl. Fig. S1E, data are shown normalized to the geometric mean of *ACT2*, *EF1α*, *GAPC2*, and *UBQ10* (Vandesompele et al. 2002). This normalization approach performed acceptably for whole seedlings, but not when roots and shoots were analyzed separately. Data are shown as the geometric mean here (Fig. S1E), because, unlike the arithmetic mean, which can be skewed by extreme outliers, the geometric mean better reflects the central tendency of multiplicative data, reducing the impact of high or low values. For the diel time course experiments using *A. thaliana*, *GAPC2* was the most consistent reference gene across all tissues and time-points (Suppl. Dataset S8). In *A. halleri*, *EF1a* was the most consistently expressed reference gene across tissues and throughout diel experiments (Suppl. Dataset S9).

### Histochemical staining and imaging

Ten freshly harvested 10-d-old seedlings cultivated on agar plates containing modified Hoagland solution, from each of seven independent homozygous lines per construct, were immersed in 1 ml of GUS staining buffer in a 2-ml polypropylene reaction vial (Zhang et al. 2022). The tubes were vacuum-infiltrated for 1 minute and then incubated in the dark at 37°C for 4 hours. The GUS staining buffer was pipetted off, and samples were cleared in 1 ml 75% (v/v) ethanol on a rocking shaker at 20 cycles per minute and RT, for 2 h, 4 times and once overnight. Seedlings were kept in 1 mL 75% (v/v) ethanol and either directly mounted on microscope slides for imaging with a light microscope (Olympus, Hamburg, Germany, Catalog no. VS120) or stored in the dark. Images shown are representative of all stained seedlings and independently transformed lines.

### Protein extraction, quantification, and β-glucuronidase activity assays

Aliquots of 100 mg frozen homogenized plant tissues were added to 200 µl of ice-cold protein extraction buffer (PEB) containing 50 mM sodium phosphate buffer (pH 7.4), 10 mM β-mercaptoethanol, 10 mM EDTA, 0.1% (v/v) SDS, and 0.1% (v/v) Triton X-100, followed by intervals of vortexing at 3,200 to 3,500 rpm for 10 s and resting on ice for 30 s until fully thawed. After centrifugation in a microcentrifuge at 4°C and 18,500×*g* for 20 minutes, the supernatant was sub-divided into several aliquots and stored at −80°C until use. Total protein quantification was performed with the Pierce™ BCA Protein Assay Kit (Thermo Fisher Scientific, Rüsselsheim am Main, Germany) using Bovine Serum Albumin (BSA) as a standard. Twenty µl of 1:10 dilutions of protein extract in a dilution buffer (50 mM sodium phosphate pH 7.4, 10 mM EDTA) were mixed with 180 µl of Bradford solution in the wells of a 96-well microtiter plate. After 10 min at RT, absorbance was measured at 595 nm in a BioTek Synergy HTX Multimode Reader plate reader (Biotek instruments, Winooski, Vermont, USA), and protein concentrations in the samples were calculated based on the standard curve. Quantification of β-glucuronidase activity was conducted as described (Jefferson et al. 1987; Gallagher 1992).

### 5’ rapid amplification of cDNA ends (RACE)-PCR

For 5’RACE analyses of of *AhHMA4*-*1* to *AhHMA4*-*3*, total RNA was extracted using Trizol from root tissues of a pool of 3 individual plants of a cross between the *A. halleri* Lan3.1 and Lan5 accessions. mRNA was then purified from total RNA using the PolyATract isolation kit (Promega, Mannheim, Germany). 5’RACE was then conducted using the purified mRNA and the SMART RACE cDNA amplification kit following the manufacturer’s instructions (Clontech, Takara Bio Europe, Saint-Germain-en-Laye, France). The PCRs were conducted using a proofreading polymerase (Strategene, Agilent Research Laboratories, Santa Clara, California, USA). The PCR products were cloned into the pGEM-T easy (Promega) and up to 10 individual clones were verified by Sanger sequencing.

### Identification and analysis of homologous promoter regions, sequence alignments, and phylogenetic analyses

Homologous segments among *HMA4* promoter regions were identified based on percentages of nucleotide identity according to Multiple Sequence Alignments of full-length promoter regions in Clustal Omega (Sievers et al. 2011). Identification and alignment of microsyntenic segments of promoter regions was done by pairwise alignment using the oline tool LALIGN and PLALIGN (https://fastademo.bioch.virginia.edu/fasta_www2/fasta_www.cgi?rm=lplalign) (Huang and Miller 1991; Madeira et al. 2019). Further processing of aligned sequences (e.g. re-alignment of misaligned positions by visual inspection, visualization, and export of high-quality images) was done in BioEdit version 5.0.9 (Hall et al. 2011). Phylogenetic analyses of promoters of *AtHMA4* and *AhHMA4-1*, *-2* and *-3* (fragments designated here full-length) were done using the Maximum Likelihood method and the Tamura-Nei model with no rate variation among sites (i.e. uniform rates) and default parameters using MEGA version 11 (Tamura and Nei 1993; Tamura et al. 2021), and bootstrapping using default parameters (Felsenstein 1985).

### Motif discovery, motif enrichment analysis, and motif comparisons

We searched for the five best motifs of between 5 and 12 nt in length with the highest statistical support occurring any number of times, in both the ER1^+^ and ER2 regions of the *AhHMA4-1, −2 and −3* promoters (as shown in Suppl. Figs. S5 and S6, Suppl. Table S3) using default options in Multiple Expectation-Maximization for Motif Elicitation (MEME) (Bailey et al. 2009; Hanikenne et al. 2013) (see Suppl. Dataset S1 and S2).

Next, we carried out multiple sequence alignments using the microsyntenic ER1^+^ and ER2 promoter segments described above. Based on MEME analyses and multiple sequence alignments, sequence motifs coinciding with MHE1 and MHE2 were identified as conserved between promoters of *AhHMA4* gene copies and across *A. halleri* accessions (Suppl. Table S3). Motif logos were generated automatically using the Download Option in the MEME suite (Bailey et al. 2009).

Next, we used the automated Motif Comparison Tool Tomtom within the MEME suite to identify transcription factor binding sites resembling MHE1 or MHE2 motifs (Gupta et al. 2007). We compared MHE1 and MHE2 motifs against three databases for plant transcription factor binding sites within the MEME suite, JASPAR2018_CORE_plants_non-redundant_v2.meme (Castro-Mondragon et al. 2022), ArabidopsisDAPv1.meme (O’Malley et al. 2016), and ArabidopsisPBM_20140210.meme(Franco-Zorrilla et al. 2014), using default parameters and the three comparison functions available in TomTom (i.e. Pearson correlation coefficient, Euclidean distance, and Sandelin-Wasserman similarity), recording each protein hit (Suppl. Dataset S3 and S4). To calculate percentages, we summed up the protein hits for each transcription factor family and divided by the total number of transcription factor protein hits. Similarly, we summed each transcription factor protein hit and divided by the total number of protein hits for that family. To assess individual occurrences of MHE1 and MHE2 motifs, and of the evening element (EE) (Harmer et al. 2000), across *HMA4* promoter regions, we used the FIMO (Find Individual Motif Occurrences) tool within the MEME suite with a *p*-value cut-off of < 10^-4^ (Grant et al. 2011) (Suppl. Dataset S5 and S6).

## Data and resource availability

Designation and sequences of the *AtHMA4* (LR782543.1, NCBI), *AlHMA4* (HE995966.1, NCBI), *AhHMA4*-*1* to *AhHMA4*-*3* (HE995813 to HE996227, EBI), the full-length promoter fragments employed (NCBI, submission id. 2881448), and *NcHMA4-1* to *-4* (HM043791.1, HM043793.1, HM043793.2, HM043793.4, NCBI) promoters are publicly available (Hanikenne et al. 2008; Lochlainn et al. 2011; Hanikenne et al. 2013). The consensus sequence of the entire *NcHMA4* genomic locus (101,480 bp) was taken from (Lochlainn et al. 2011). The *AtHMA4* promoter regions of 1,135 *A. thaliana* accessions were extracted using the 1001 *A. thaliana* genome tools (Alonso-Blanco et al. 2016) accessed *via* the Pseudogenomes Download option by specifying the chromosomal region of interest (i.e. Chr2:8276875..8279477; https://1001genomes.org/). Megablast searches (Boratyn et al. 2013) using full-length *AtHMA4_P_*, *AlHMA4_P_*, and *AhHMA4*-*1_P_*, *4-2_P_* and *4-3_P_* sequences as queries were used for retrieving similar *HMA4* promoter sequences from other land plant species (taxid: 3193). We selected the blast hits with a query coverage of ≥ 25% and obtained *HMA4* promoter sequences from *Arabis alpina* (LT669790.1, NCBI), *Arabidopsis arenosa* (LR999453.1, NCBI), *Capsella bursa-pastoris* (CP144185.1, NCBI), *Camelina sativa* (CP146050.1, NCBI), *Brassica napus* (CP151901.1, NCBI), *B. oleracea* (LR031876.1, NCBI), *B. rapa* (LR031574.1, NCBI), *Raphanus sativus* (LR778310.1, NCBI), and *Thlaspi arvense* (OU466859.2, NCBI). Further data underlying this article are available in the article and in its online supplementary material. Any other data and materials will be shared upon reasonable request to the corresponding author.

## Author contributions

LC, MH and UK conceived the project; LC and UK designed research; all shown experiments were conducted by LC; MH, JC, CN, and JS generated constructs for the initial deletion series. LC and JC generated all follow-up constructs based on *AhHMA4-1_P_* and *AtHMA4_P_*. LC and UK analyzed data; LC and UK wrote the manuscript. All authors read and edited the manuscript.

## Acknowledgments

We are grateful to Andreas Aufermann and Martin Pullack for technical assistance in plant cultivation and all lab members of the Ruhr University Bochum, Germany, for comments. We also thank Dr. Hassan Ahmadi for technical assistance in hydrophonic cultivation of *A. halleri* and *A. thaliana* plants. This work was funded by the German Research Foundation (Deutsche Forschungsgemeinschaft, DFG) grant 1967/3-3 to U.K., by the National Council of Humanities, Sciences and Technologies (Consejo Nacional de Humanidades, Ciencias y Tecnologías, CONACYT) scholarship no. 438349, by ERC-AdG 788380 “LEAP EXTREME” to U.K., by Ruhr University Bochum, Bochum, Germany, as well as “Fonds de la Recherche Scientifique–FNRS” (FRFC-2.4583.08 and PDR-T.0206.13, to M.H.), the University of Liège (SFRD-12/03) (M.H.), and the Belgian Program on Interuniversity Attraction Poles (IAP no. P7/44) (M.H.).

## Conflict of Interest Statement

The authors declare no conflict of interest.

## Notes

### Competing Interest Statement

The authors have declared no competing interest.

