## Supplementary Figures and Tables for "*Cis*-regulatory mutations co-opting circadian clock regulation underlie naturally selected extreme trait in *Arabidopsis halleri*"

Supplementary Figure S1. Sequence and functional conservation among *A. halleri* and *A. thaliana HMA4* promoter regions

Supplementary Figure S2. Quantitative relationships between *GUS* reporter transcript levels and specific GUS activity for *HMA4* promoters and deletion series

Supplementary Figure S3. Functions and annotations of the distal regions of *HMA4* promoters of *A. thaliana* and *A. halleri*

Supplementary Figure S4. Dissection of ER1 in the *AhHMA4-1* promoter and identification of the candidate *cis*-regulatory enhancer element MHE1

Supplementary Figure S5. Conservation of the ER1$^{+}$ with candidate *cis*-regulatory enhancer element MHE1 across the promoters of the three *HMA4* gene copies of multiple *A. halleri* genotypes

Supplementary Figure S6. Conservation of the ER2 with two candidate *cis*-regulatory enhancer elements MHE2 across the promoters of the three HMA4 gene copies of multiple *A. halleri* genotypes

Supplementary Figure S7. Reproducibility of the enhancement and diel dynamics of GUS reporter transcript levels dependent on MHE2 and CCA1 in *A. thaliana*

Supplementary Figure S8. Reproducibility of diel dynamics of *HMA4* transcript levels in *A. halleri*

### Supplementary Tables

Supplementary Table S1. Primers used in this study

Supplementary Table S2. Cloning procedures, primers, and plasmids used in this study

Supplementary Table S3. Motifs identified in ER1$^{+}$ and ER2 segments of *AhHMA4* promoters of *A. halleri* accessions

**A**

| | *AhHMA4-1*$_P$ | *AhHMA4-2*$_P$ | *AhHMA4-3*$_P$ |
|---|---|---|---|
| ***AtHMA4*$_P$** | 78 | 49 | 46 |
| ***AhHMA4-1*$_P$** | | 52 | 51 |
| ***AhHMA4-2*$_P$** | | | 93 |

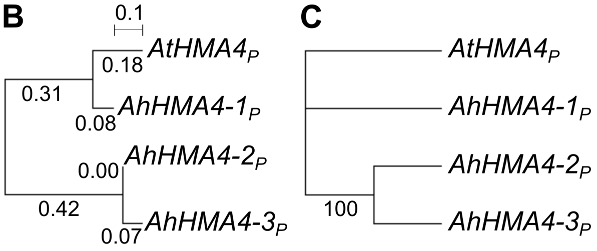


**D**

<table>
<tr><th></th><th colspan="5">AhHMA4-1$_P$</th><th colspan="8">AhHMA4-2$_P$</th><th colspan="9">AhHMA4-3$_P$</th></tr>
<tr><th></th><th>DR</th><th>IR</th><th>PR</th><th>5'UTR</th><th>i</th><th colspan="2">DR</th><th>IR</th><th colspan="2">PR</th><th>5'UTR</th><th colspan="2">i</th><th colspan="2">DR</th><th>IR</th><th colspan="2">PR</th><th colspan="2">5'UTR</th><th colspan="2">i</th></tr>
<tr><td>AtHMA4$_P$</td><td>77[a]</td><td>70[a]</td><td>85[a]</td><td>72[a]</td><td>84[a]</td><td colspan="2">53[c]</td><td>50[c]</td><td colspan="2">73[c]</td><td>61[a]</td><td>75[a]</td><td>64[a]</td><td>72[c]</td><td>53[c]</td><td>47[c]</td><td colspan="2">62[c]</td><td>57[b]</td><td>65[a]</td><td>75[a]</td><td>67[a]</td></tr>
<tr><td>AhHMA4-1$_P$</td><td colspan="5"></td><td>53[c]</td><td>82[c]</td><td>56[d]</td><td>80[c]</td><td>78[c]</td><td>83[a]</td><td>81[b]</td><td>62[a]</td><td>52[c]</td><td>74[c]</td><td>62[d]</td><td colspan="2">61[c]</td><td>56[b]</td><td>83[a]</td><td>81[a]</td><td>62[a]</td></tr>
<tr><td>AhHMA4-2$_P$</td><td colspan="13"></td><td colspan="2">96[a]</td><td>98[a]</td><td>74[b]</td><td>92[a]</td><td colspan="2">86[a]</td><td colspan="2">95[a]</td></tr>
</table>

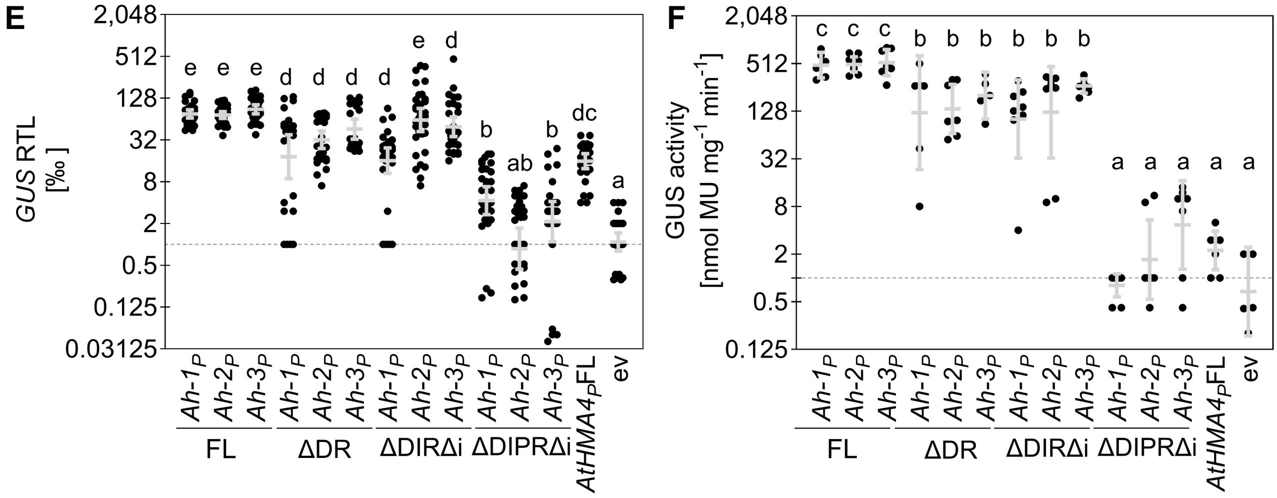


**Supplementary Figure S1. Sequence and functional conservation among *A. halleri* and *A. thaliana HMA4* promoter regions.**
**(A)** Global comparisons between full-length promoters of *AhHMA4-1* (1,602 nt), *-2* (1,761 nt), and *-3* (1,494 nt) and *AtHMA4* (2,000 nt) promoters, based on multiple sequence alignments using Clustal OMEGA. Numbers are percentages of identical nucleotides. The full alignment (3,050 nt) is shown in Supplementary Dataset S11. **(B and C)** Maximum likelihood trees of full-length *HMA4* promoter sequences, including the tree with the highest log likelihood (-8795, B) and a bootstrapped consensus tree inferred from 1,000 replicates using the Maximum Likelihood method and Tamura-Nei model in MEGA11 (C). The 3,050 nt long global sequence alignment between *AhHMA4-1*$_P$, *-2*$_P$, and *-3*$_P$ and *AtHMA4*$_P$ (see A) was used to build the tree using MEGA11, omitting positions containing gaps and missing data. Initial tree(s) for the heuristic search were obtained automatically by applying Neighbor-Join and BioNJ algorithms to a matrix of pairwise distances estimated using the Tamura-Nei model, and then automatically selecting the topology with maximal log likelihood value. The scale bar represents the distance scale, and numbers on branches indicate substitutions per site (B).
The percentage of replicate trees in which the associated taxa were grouped together in the bootstrap test are shown next to the respective branches, and branches corresponding to partitions reproduced in less than 50% bootstrap replicates are collapsed (C).
**(D)** Comparisons among sequence segments constituting *HMA4* promoter subregions (see Fig. 1A). Numbers are percentages of identical nucleotides. Superscript characters indicate *e*-value of < 0.0001[a], < 0.01[b], < 1[c], and ≤ 100[d] based on pairwise-alignments using the LALIGN/PLALIGN local alignment tool (see Methods). **(E and F)** Relative *GUS* transcript levels (E) and specific GUS enzyme activity (F) in whole seedlings of *A. thaliana GUS* reporter lines for full-length and 5' deletion series of the *HMA4-1* (*Ah-1*$_P$), *HMA4-2* (*Ah-2*$_P$) and *HMA4-3* (*Ah-3*$_P$) promoters of *A. halleri*. *GUS* transcript levels are shown normalized to the geometric mean of those of *ACT2*, *EF1α*, *GAPC2*, and *UBQ10* (Vandesompele et al. 2002) and subsequently multiplied by one thousand (E). Bars show mean ± SD ($n$ = 5 to 33 datapoints) of independent transgenic lines, with each datapoint representing the mean of three replicate multi-well plates of qPCR reactions (E), or multi-well enzyme assays (F). Different characters indicate statistically significant differences (one-way non-parametric ANOVA with Dunn´s multiple comparison test, $P$ < 0.05). For the definition of abbreviations see Figs. 1 and 2.

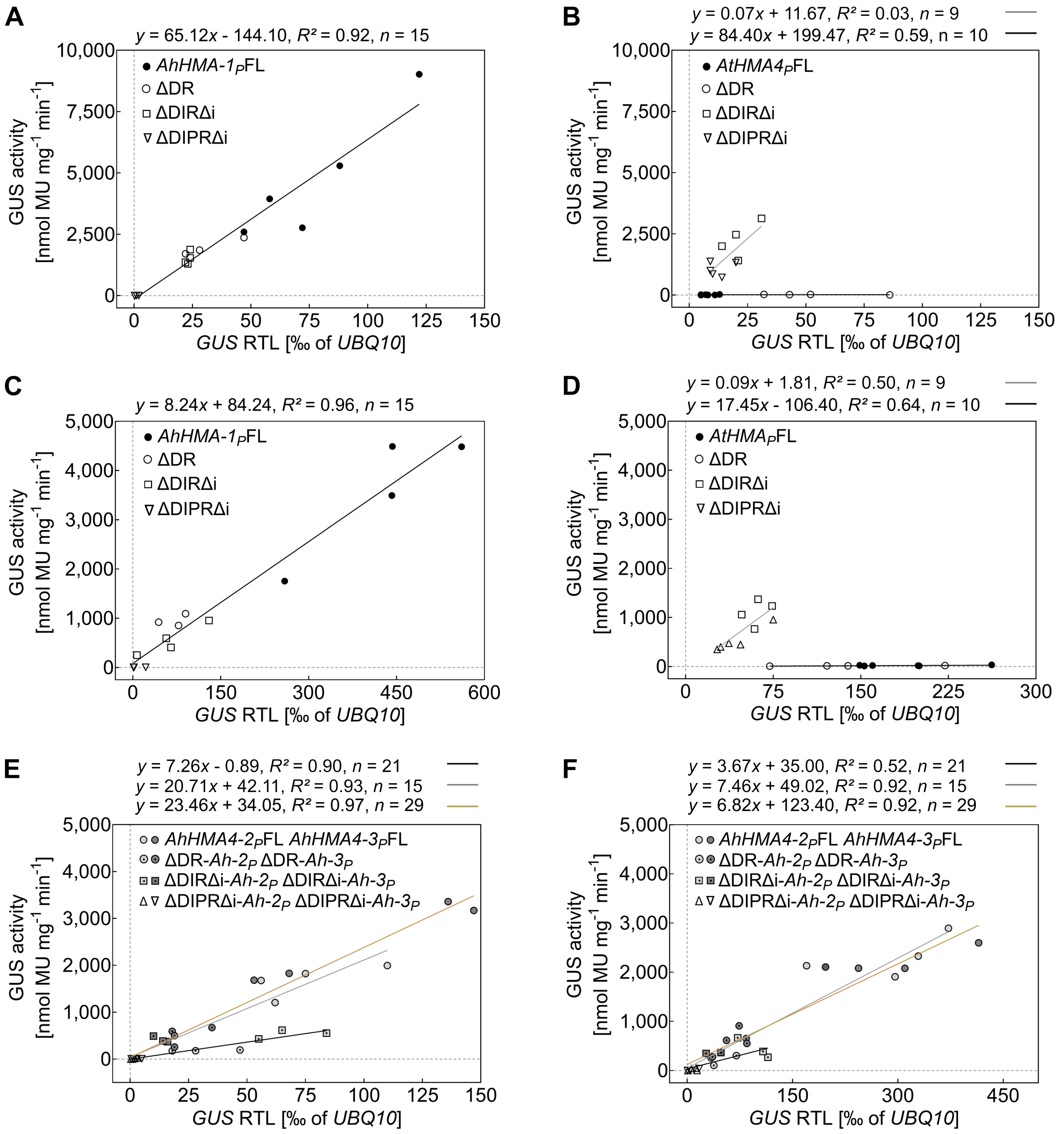


**Supplementary Figure S2. Quantitative relationships between *GUS* reporter transcript levels and specific GUS activity for *HMA4* promoters and deletion series. (A-F)** Plots showing datapoints and linear regressions for specific GUS enzyme activities as a function of *GUS* transcript levels in shoots (A, B, E) and roots (C, D, F) of *A. thaliana GUS* reporter lines for *AhHMA4-1*$_P$ (A and C), *AtHMA4*$_P$ (B and D), and *AhHMA4-2*$_P$ and *-3*$_P$ (E and F) including the data for all constructs shown in Fig. 2 and part of the constructs shown in Fig. 1. Equations and coefficients of determination ($R^2$) are shown above each diagram. For the definition of abbreviations and the designations of constructs see Figs. 1 and 2.

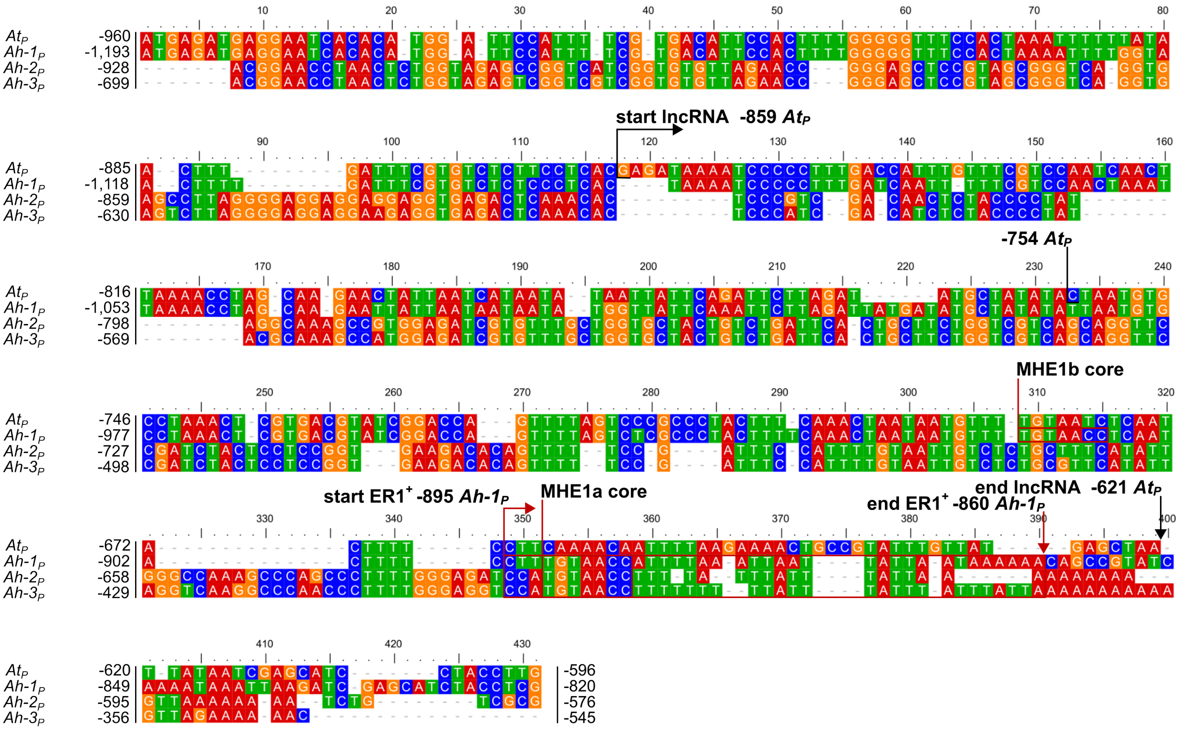

**Supplementary Figure S3. Functions and annotations of the distal regions of *HMA4* promoters of *A. thaliana* and *A. halleri*.** Multiple sequence alignment of part of the distal regions of the promoters of *AtHMA4* and *AhHMA4-1*, *4-2*, and *4-3*. Fragments corresponding to the distal region of 363 nt of *AtHMA4P* ($At_P$), 374 nt of $AhHMA4\text{-}1_P$ ($Ah\text{-}1_P$), 353 nt of $AhHMA4\text{-}2_P$ ($Ah\text{-}2_P$), and 355 nt of $AhHMA4\text{-}3_P$ ($Ah\text{-}3_P$), were used to generate the alignment using the ClustalW algorithm with default options in MEGA v.11.0, followed by processing in BioEdit v.5.0.9 (see Methods). The numbers on the left indicate positions relative to the respective transcriptional start site. The annotation of the lncRNA (AT2G07115.1) was taken from TAIR (The Arabidopsis Information Resource). The start of the annotated lncRNA in $AtHMA4_P$ is indicated by a black cornered arrow above the alignment, and the end by a black downward vertical arrow below the alignment. A sequence with elevated degree of sequence similarity among the promoters of the three *A. halleri HMA4* gene copies (-895 to -860 in $AhHMA4\text{-}1_P$), addressed here as enhancing subregion 1 ($ER1^+$), is indicated by a red cornered arrow (start) and by a red downward vertical arrow (end). The TGT AAC core Metal Hyperaccumulation Elements 1a and 1b (MHE1a, MHE1b) is boxed in red in *AhHMA4* promoters and additionally highlighted by red horizontal lines above the alignment between positions -914 to -908 and -892 to -886 of $AhHMA4\text{-}1_P$ (see also Fig. 3, Suppl. Fig. 4).

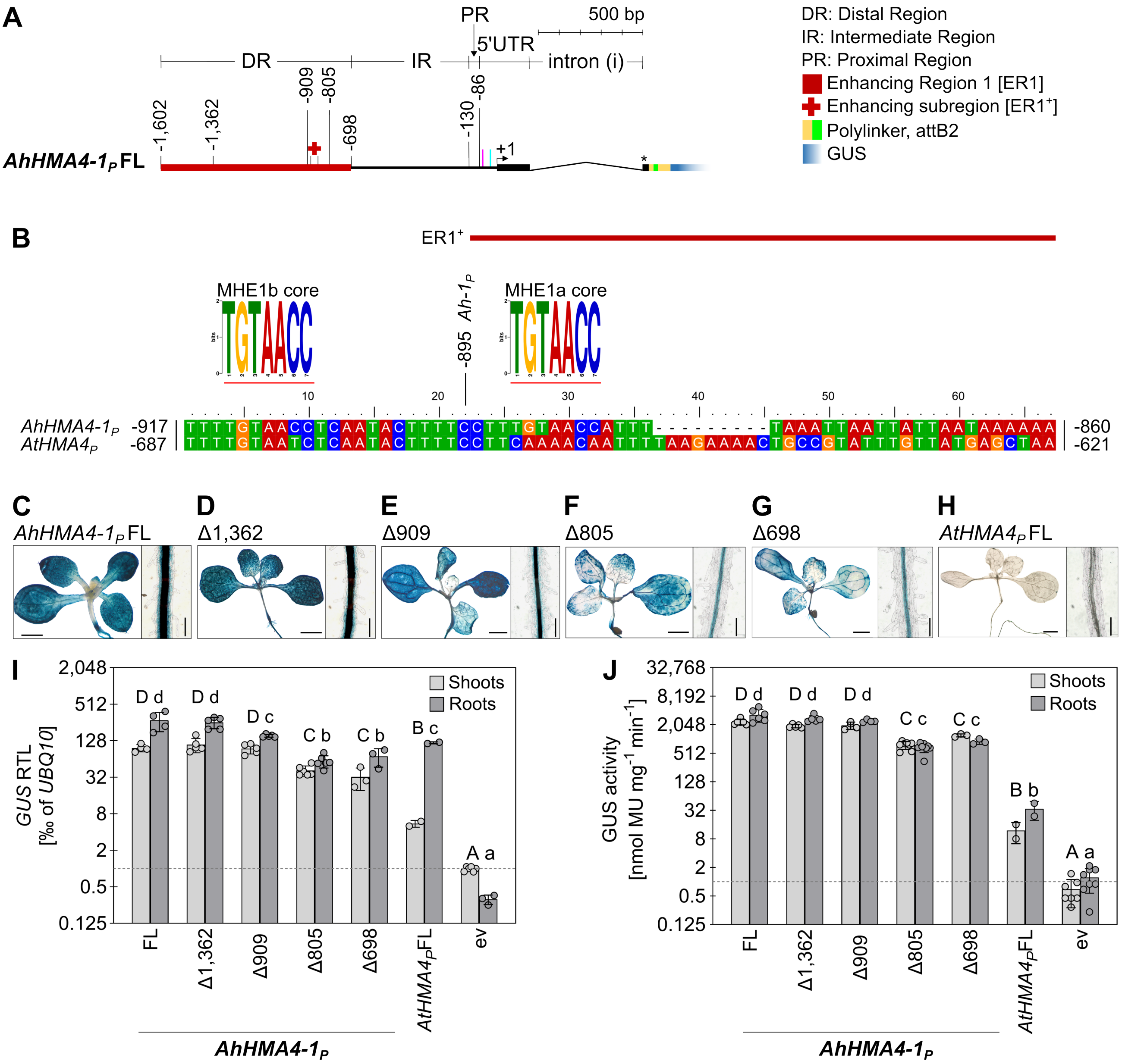


**Supplementary Figure S4. Dissection of ER1 in the *AhHMA4-1* promoter and identification of the candidate *cis*-regulatory enhancer element MHE1. (A)** Scheme of the *AhHMA4-1* promoter construct to outline the partial deletions of Enhancing Region 1 (ER1) to identify ER1$^+$. Numbers indicate the distance in bp from the transcriptional start site (+1), for the breakpoints implemented in the promoter deletion series of reporter constructs introduced into *A. thaliana*. Positions of the CAAT/TATA box are marked by magenta/cyan vertical lines. **(B)** Alignment of the ER1$^+$ region of *AhHMA4-1*$_P$ (nucleotides -895 to -860), including the upstream 21 nt region (nucleotides -917 to -896) that contains a second copy of the core MHE1 motif (MHE1b, see Fig. 3B), with the homologous region of *AtHMA4*$_P$. The logos of the core MHE1 motifs (underlined in red) are shown above the corresponding positions in the alignment, with MHE1a within ER1$^+$ (marked by a horizontal dark red line above) and MHE1b 22 nt upstream (MHE1b is missing in *AhHMA4-2*$_P$ and *AhHMA4-3*$_P$, see Suppl. Fig. S3). **(C-H)** Histochemical detection of GUS activity in rosettes (left) and the root hair zone of roots (right) for full-length *AhHMA4-1*$_P$ (C), and *AtHMA4*$_P$ (H), and partial deletions of ER1 (D to G). Size bars: 1 mm (rosettes), and 0.1 mm (roots). **(I-J)** Relative *GUS* transcript levels and specific GUS enzyme activities in transgenic *A. thaliana GUS* reporter lines carrying partial deletions of ER1. Bars show mean ± SD ($n$ = 3 to 9) of independent transgenic lines. Each datapoint represents the mean of three technical replicate multi-well microplates of qPCR (I) or enzyme assays (J). Two independent transgenic lines carrying full-length *AtHMA4*$_P$ (FL) were included here. Different characters indicate statistically significant differences based on one-way non-parametric ANOVA followed by a Dunn's multiple comparison test ($P$ < 0.05). ev, transformants with an empty vector (see Fig. 1A).

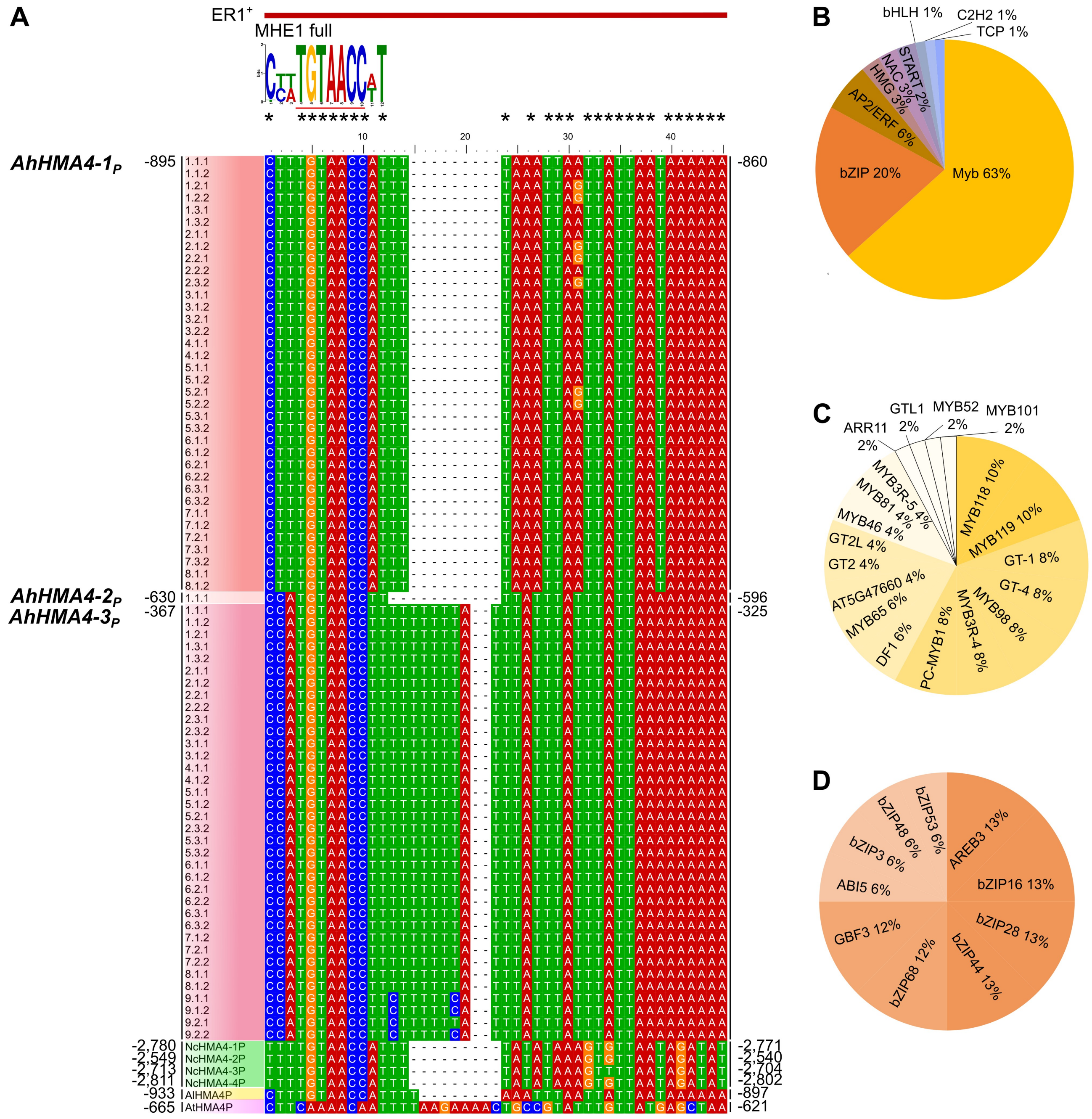


**Supplementary Figure S5. Conservation of the ER1$^+$ with candidate *cis*-regulatory enhancer element MHE1 across the promoters of the three *HMA4* gene copies of multiple *A. halleri* genotypes. (A)** Multiple sequence alignment of enhancing sub-region ER1$^+$, with the sequence logo of the Metal Hyperaccumulation Element 1a (MHE1a) motif shared by the promoters of all three *AhHMA4* gene copies in multiple *A. halleri* individuals from different geographic origins (corresponding to MHE1a, see Suppl. Fig. S4; data from Hanikenne et al. 2013). ER1$^+$ is marked by a dark red line above the alignment. The MHE1 core motif is underlined in red. Microsyntenic *HMA4* promoter regions of the hyperaccumulator *Noccaea caerulescens* (green), *A. lyrata* (yellow), and *A. thaliana* (pink) are included at the bottom. Numbers identifying sequences specify the *A. halleri* population (i.e. collection site), individual, and allele (Hanikenne et al. 2013). Note that only a single ER1$^+$ sequence is available for *AhHMA4-2$_P$* (from Lan3.1 accession). **(B-D)** Percentage representation of protein families ($n$ = 9) (B) among transcription factor proteins predicted to bind MHE1 (redundant, $n$ = 24; non-redundant, $n$ = 13; total $n$ = 82), and of specific transcription factors (C and D) belonging to the three most frequently predicted protein families (see B). Representative frequently predicted transcription factors are in the MYB (INTERPRO ID: IPR006447) (C), and bZIP (INTERPRO ID: PR004827) (D) protein families (full list in Suppl. Dataset S3). The motif shown in (A) was used for generating predictions applying three motif similarity comparison functions (Pearson correlation coefficient, Euclidean distance, and Sandelin-Wasserman similarity) across three databases containing plant transcription factor binding sites (JASPAR CORE 2022 plants by Castro-Mondragon et al. 2022, DAP motifs by O'Malley et al. 2016, and PBM motifs by Franco-Zorrilla et al. 2014). Both the motif similarity comparison functions and databases are integrated into the Tomtom tool in the MEME suite. Each protein family or transcription factor protein predicted in each database by was interpreted as one occurrrence.

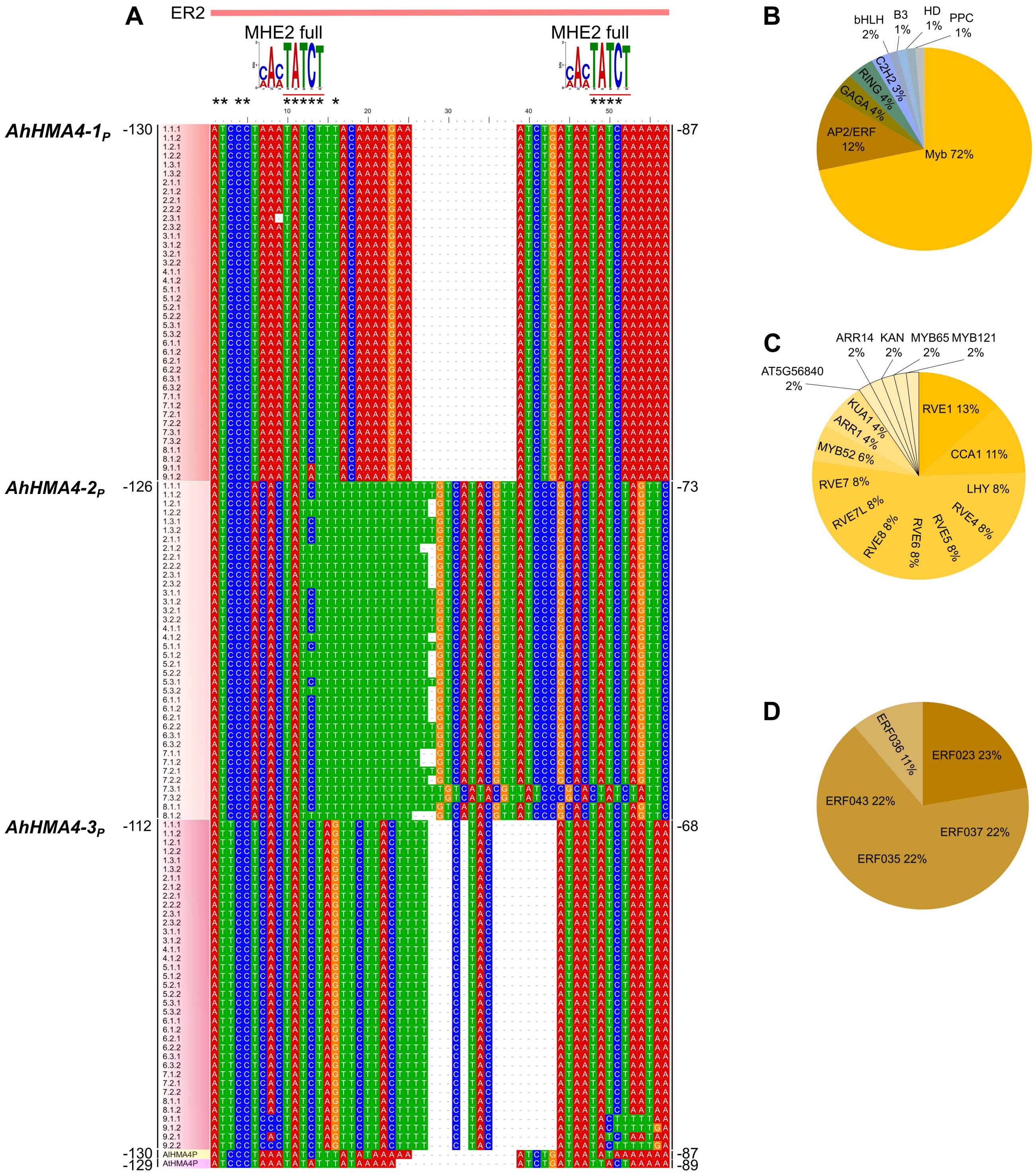


**Supplementary Figure S6. Conservation of the ER2 with two candidate *cis*-regulatory enhancer elements MHE2 across the promoters of the three *HMA4* gene copies of multiple *A. halleri* genotypes. (A)** Multiple sequence alignment of enhancing subregion (ER2), with the sequence logo of Metal Hyperaccumulation Element 2 (MHE2) motif shared by the promoters of all three *AhHMA4* gene copies in multiple *A. halleri* individuals from different geographic origins (corresponding to MHE2a and b, see Fig. 3C; data from Hanikenne et al. 2013). ER2 is marked by a light red line above the alignment. The MHE2 core motif is underlined in red. Microsyntenic *HMA4* promoter regions of *A. lyrata* (yellow), and *A. thaliana* (pink) are included at the bottom. Numbers identifying sequences specify the *A. halleri* population (i.e. collection site), individual, and allele (see Hanikenne et al. 2013). **(B-D)** Percentage representation of protein families ($n$ = 9) (B) among transcription factors predicted to bind MHE2 (redundant, $n$ = 18; non-redundant, $n$ = 15; total $n$ = 74), and of specific transcription factors belonging to the frequently identified families predicted to bind MHE2 (MYB-related in C, AP2/ERF in D, see B; full list in Suppl. Dataset S4). See also Suppl. Fig. S5.

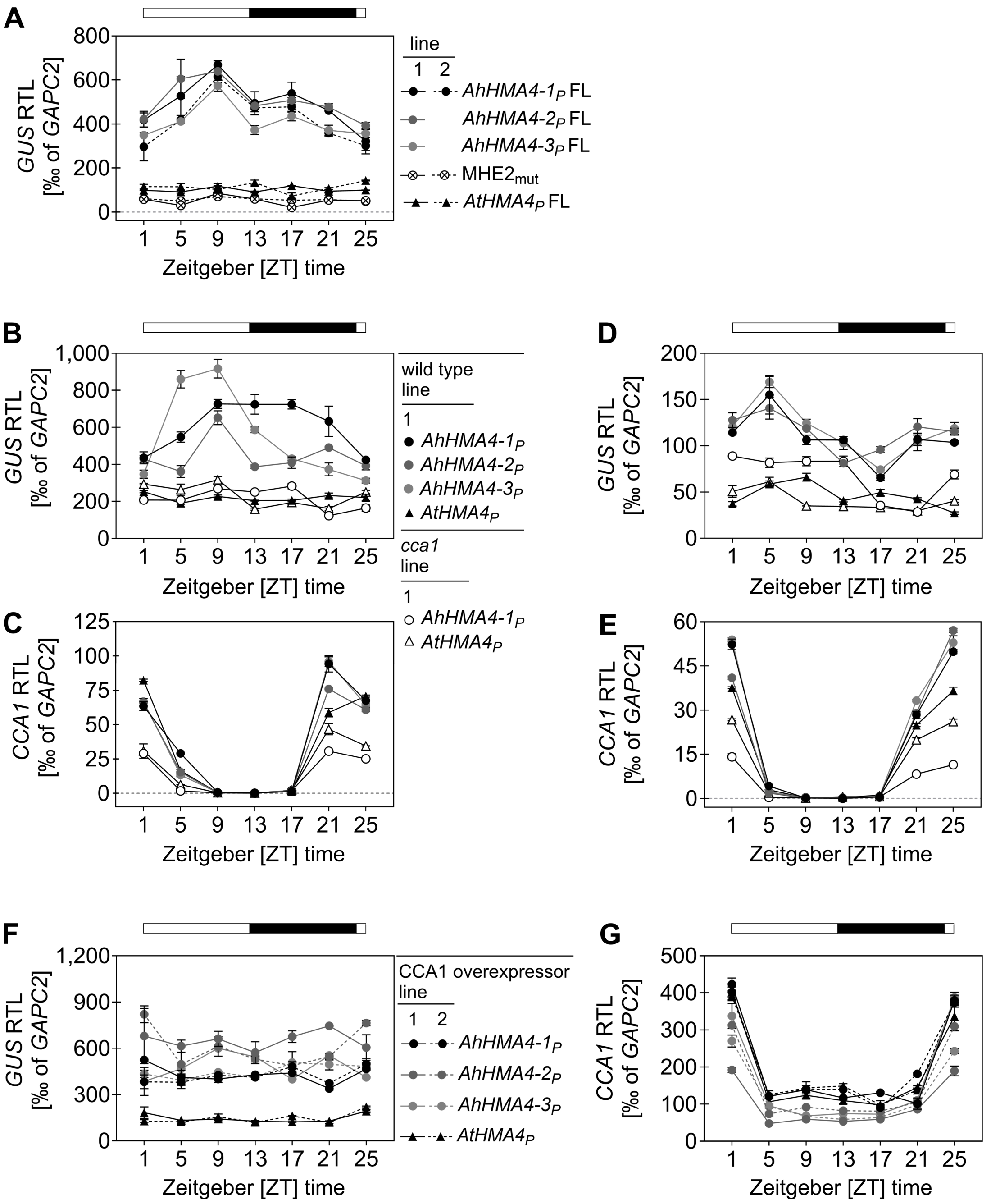


**Supplementary Figure S7. Reproducibility of the enhancement and diel dynamics of *GUS* reporter transcript levels dependent on MHE2 and *CCA1* in A. thaliana. (A)** Relative *GUS* transcript levels quantified over a diurnal cycle in seedlings of transgenic *A. thaliana* promoter-*GUS* reporter lines (independent experiment, see Fig. 5A). Datapoints are mean ± SD (*n* = 3 technical replicates as independent qPCR plates). Two independent transgenic lines (1, 2) are shown per construct, and one representative line for *AhHMA4-2*$_P$ and *4-3*$_P$. Seedlings were cultivated on agar plates and harvested on day 21. *GUS* reporter lines contain the full-length (FL) *AhHMA4-1*$_P$, *4-2*$_P$, *4-3*$_P$, the *AtHMA4*$_P$ FL, or the *AhHMA4-1*$_P$ FL construct with both copies of MHE2 mutated ($MHE2_{mut}$) by site-directed mutagenesis to match the corresponding sequence of *AtHMA4*$_P$ (see also Fig. 3A and 3C). **(B-E)** Relative *GUS* and *CCA1* transcript levels in shoots (B and C), and roots (D and E) of wild-type and *cca1-1* mutant *A. thaliana* transgenic promoter-*GUS* reporter lines as indicated in (B). Datapoints are mean ± SD (*n* = 3 technical replicates as independent qPCR plates) from one representative experiment. Promoter-*GUS* transformant lines were crossed with a *cca1* mutant plant, and the experiment was done using homozygous plants of the F3 generation. **(F and G)** Relative *GUS* (F) and *CCA1* (G) transcript levels in seedlings of the wild-type and a *CCA1* overexpressor line of *A. thaliana* transgenic promoter-*GUS* reporter lines as indicated in (F). Two independent promoter-*GUS* transformant lines (1, 2) are shown per construct, each of which was crossed with the *CCA1* overexpressor line, and the experiment was done using homozygous plants of the F3 generation). Horizontal bar: white fill, day; black fill, night (A-G).

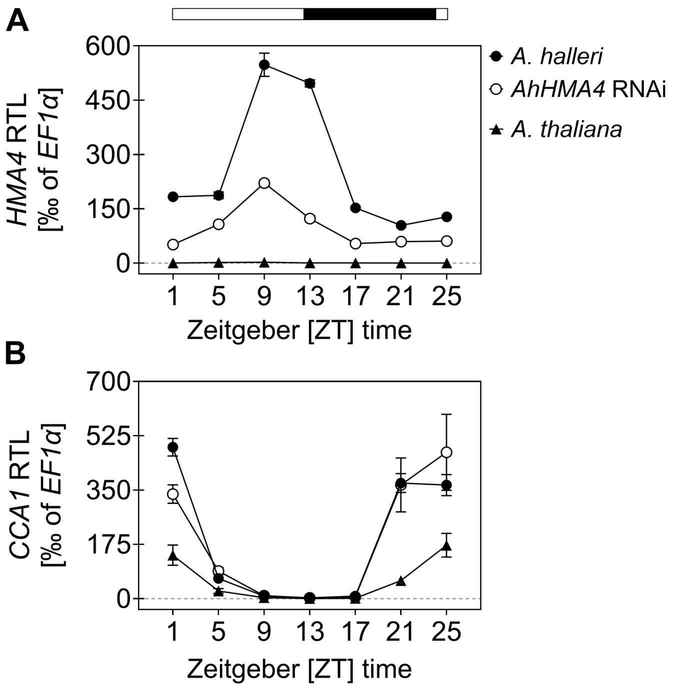


**Supplementary Figure S8. Reproducibility of diel dynamics of *HMA4* transcript levels in *A. halleri*. (A-D)** Diel time course of relative *HMA4* and *CCA1* transcript levels in shoots (A and B) and roots (C and D) of *A. halleri* wild-type plants and a *HMA4* RNAi line, as well as in *A. thaliana*. Datapoints are mean ± SD ($n$ = 3 technical replicates as independent qPCR plates, from a second representative experiment of a total of two experiments, see Fig. 5E-H). Plants were cultivated in hydroponics and harvested on day 21. Horizontal bar: white fill, day; black fill, night.

**Supplementary Table S1. Primers used in this study**

| (A) Primers used for cloning | Sequence (5´to 3´) |
|---|---|
| FP#1 | GGCCGCCCCCTTCACCAATTAACTTGGTTTCAAGACTAATTAAGATCG |
| RP#1 | CGATCTTAATTAGTCTTGAAACCAAGTTAATTGGTGAAGGGGGCG GCC |
| FP#2 | GGCCGCCCCCTTCACCAAAGATAGCCTACGAGATTTACGTGG |
| RP#2 | CCACGTAAATCTCGTAGGCTATCTTTGGTGAAGGGGGCGGCC |
| FP#3 | GGCCGCCCCCTTCACCGGTTTAATAAAGGTTTCAAAAATTTAG |
| RP#3 | CTAAATTTTTGAAACCTTTATTAAACCGGTGAAGGGGGCGGCC |
| FP#4 | GGCCGCCCCCTTCACCGGTATTTTTAATAATTTTGAATTTTTGTGG |
| RP#4 | CCACAAAAATTCAAAATTATTAAAAATACCGGTGAAGGGGGCGGC C |
| FP#5 | GGCCGCCCCCTTCACCATCCCTAAATATATTTATAAAAAATCTG |
| RP#5 | CAGATTTTTTATAAATATATTTAGGGATGGTGAAGGGGGCGGCC |
| FP#6 | CTAGCTCTCCTCTCTTCTCCGAAAATGGCGTTACAAAACAAAG |
| RP#6 | CTTTGTTTTGTAACGCCATTTTCGGAGAAGAGAGGAGAGCTAG |
| FP#7 | GGCCGCCCCCTTCACCAAAAACTTAGTTCAAATTTTAAGAAGTTC |
| RP#7 | GAACTTCTTAAAATTTGAACTAAGTTTTTGGTGAAGGGGGCGGCC |
| FP#8 | GGCCGCCCCCTTCACCAAAAATAGCCTACGAAATTTACATGG |
| RP#8 | CCATGTAAATTTCGTAGGCTATTTTTGGTGAAGGGGGCGGCC |
| FP#9 | GGCCGCCCCCTTCACCATCCCTAAATATCTTTACAAAAGAAATCTG |
| RP#9 | CAGATTTCTTTTGTAAAGATATTTAGGGATGGTGAAGGGGGCGGCC |
| FP#10 | CTAGCTCTCCTCTCTTCTCCGAAAATGGCGTTACAAAACAAAG |
| RP#10 | CTTTGTTTTGTAACGCCATTTTCGGAGAAGAGAGGAGAGCTAG |
| FP#11 | GGCCGCCCCCTTCACCTATTTAATAGGTAGTGCATCATATTCG |
| RP#11 | CGAATATGATGCACTACCTATTAAATAGGTGAAGGGGGCGGCC |
| FP#12 | TAGCTTGCGTCTCTTCTCCGAAAATGGCGTCACAAAACAAAG |
| RP#12 | CTTTGTTTTGTGACGCCATTTTCGGAGAAGAGACGCAAGCTAG |
| FP#13 | GGCCGCCCCCTTCACCCGTTATCCCACACTATCTTTTTTTTTTTTTGTC |
| RP#13 | GACAAAAAAAAAAAAAGATAGTGTGGGATAACGGGTGAAGGGGGCGGCC |
| FP#14 | CTAGTTAATTAACATTTTTTTTTTCCTATTTATCTGCC |
| RP#14 | CTACAGGCATCGTGGTGTCAC |
| FP#15 | GGCAGATAAATAGGAAAAAAAAAATGTTAATTAACTAGAGCTTGGCAC |
| RP#15 | GTGACACCACGATGCCTGTAG |
| FP#16 | GTGCCAAGCTCTAGTTAATTAACTGTCGCGATACTTACCAAACAAATC |
| RP#16 | GATCCTCTAGAGTCGAGGCGCGCC |
| FP#17 | CTAGTTAATTAACATACGTTATTCCTCACTATC |
| FP#18 | GATAGTGAGGAATAACGTATGTTAATTAACTAGAGCTTGGCAC |
| FP#19 | CTAGTTAATTAACATAATATCTAATAAATATCTAG |
| FP#20 | CTAGATATTTATTAGATATTATGTTAATTAACTAGAGCTTGGCAC |
| FP#21 | *CACC*CTAATGTGCCTAAACTCGTGACG |
| RP#17 | GGGAGAGATGAGGAGACTGTCGTG |
| FP#22 | *CACC*AATTAACTTGGTTTCAAGACTAATTAAGATCG |
| FP#23 | GTTTTGTAACCTCAATACTTTTCCTTC |
| RP#18 | GTATTGAGGTTACAAAACATTATTAGTTTGAAAGTAGG |
| FP#24 | CCTTTGTAACCATTTTAAGAAAACTGC |
| RP#19 | CTTAAAATGGTTACAAAGGAAAAGTATTGAG |
| FP#25 | CCTAAATATCTTTACAAAAAATCTGATAATATCAAAAATGTTAAAG |
| RP#20 | CATTTTTGATATTATCAGATTTTTTGTAAAGATATTTAGGGATAATG |
| FP#26 | *CACC*ATCCCTAAATATATTTATAAAAAATC |
| FP#27 | *CACC*TGTTAAAGAAAACTAACC |
| FP#28 | *CACC*CGGGTTAATATCTAGTTAGAAAG |
| RP#21 | TTTCTCTTCTTCTTTGTTTTGTGACGC |
| FP#29 | *CACC*CCTCAATACTTTTCCTTTGTAACC |
| FP#30 | *CACC*CAACTTTCAAGACTAATTAAGATCGAG |
| FP#31 | *CACC*TGTTAAAGAAAACCAACCAATCATCTTGC |
| FP#32 | CACGGGACTGGTTATATTTCGG |
| RP#22 | CTTTAACAGAACTATAAAATATTATAGGGTAATTCG |
| FP#33 | CCTATAATATTTTATAGTTCTGTTAAAGAAAACC |
| FP#34 | *CACC*CACGGGACTGGTTATATTTCGG |
| FP#35 | *CACC*ATCCCTAAATATCTTTACAAAAG |
| FP#36 | *CACC*TGTTAAAGAAAACCAACCAATCATCTTGC |
| FP#37 | GTTTCAAAACATCAATACTTTTCCTTTG |
| RP#23 | GAAAAGTATTGATGTTTTGAAACATTATTAG |
| FP#38 | CCTTCAAAACAATTTTAAATTAATTATTAATAAAAAACAGCCGTATC |
| RP#24 | GCTGTTTTTTATTAATAATTAATTTAAAATTGTTTTGAAGGAAAAGTATTG |
| FP#39 | CCTAAATATATTTATAAAATCAATCTGATAATTACTAAAAATGTTAAAG |
| RP#25 | CATTTTTAGTAATTATCAGATTGATTTTATAAATATATTTAGGGATAATG |
| FP#40 | *CACC*ACTTACCGATCGGGTATGCCATG |
| RP#26 | TTTCTGTAAAACGTAAATCAAAACAAC |
| FP#41 | CACGGGACTGGTTATATTTCGG |
| RP#27 | GTCTTGAAACCAAGTTAATTTTTTTTTTAATACAAATTGTCATAACTCATAAG |
| FP#42 | CTTATGAGTTATGACAATTTGTATTAAAAAAAAAATTAACTTGGTTTCAAGACTAATTAAGATCG |
| RP#28 | AC |
| FP#43 | GGGAGAGATGAGGAGACTGTCGTG |
| | *CACC*CACGGGACTGGTTATATTTCGG |

*Continued on the next page*

| **(B) Primers used for RT-qPCR** | **Sequence (5´to 3´)** |
|---|---|
| FP-*AtEF1α* (AT5G60390)[1]* | TGAGCACGCTCTTCTTGCTTTCA |
| RP-*AtEF1α* (AT5G60390)[1]* | GGTGGTGGCATCCATCTTGTTACA |
| FP-*AtACT2* (AT3G18780)[2] | TCTCTCTTCCTCATGCCA |
| RP-*AtACT2* (AT3G18780)[2] | TCTGTAAGGATCTTCATGAGGTAAT |
| FP-*At*GAPC2 (AT1G13440)[2]* | TAACTGCCTTGCTCCTCTT |
| RP-*At*GAPC2 (AT1G13440)[2]* | AGAGTGGACAGTGGTCA |
| FP-*AtUBQ10* (AT4G05320)* | GGCCTTGTATAATCCCTGATGAATAAG |
| RP-*AtUBQ10* (AT4G05320)* | AAAGAGATAACAGGAACGGAAACATAGT |
| FP-*GUS* (*uidA*) | ACAAAAACCACCCAAGCGTG |
| RP-*GUS* (*uidA*) | GCCAGTGGCGCGAAATATT |
| FP-*AtCCA1* (AT2G46830) | GAGGAGAAGAGAAACACAGG |
| RP-*AtCCA1* (AT2G46830) | CTACTCATTAGCTTTGAAGCAT |
| FP-*AhCCA1* | GAGGAGAGGAGAAACACAGG |
| RP-*AhCCA1* | CTTATTAATAGCTTTGAAGCAT |
| FP-*AtHMA4* (AT2G19110) | GCTACAGCTGATATTGGTATCT |
| RP-*AtHMA4* (AT2G19110) | ATTACCAGTTTGTGTTGCAAG |
| FP-*AhHMA4* | CCCTCTCTCAACCTTTATGGTAACAC |
| RP-*AhHMA4* | GCTAGTAGCAAAAGGAAGAAGCCG |

| **(C) Primers used for genotyping** | **Sequence (5´to 3´)** |
|---|---|
| LB-*cca1-1*[3] | GATGCACTCGAAATCAGCCAATTTTAGAC[a] |
| FP-*cca1-1*[4] | TGAGATTTCTCCATTTCCGTAGCTTCTGG[a] |
| RP-*cca1-1*[4] | ATCCGTTTGGGATCTTTCTGTTCCACATG[a] |
| FP-*GUS*[b] | CCGGCGGGATAGTCTGCC |
| RP-*GUS*[b] | GCCAGTGGCGCGAAATATT |

[1]Hanikenne et al. 2008, [2]Lai et al. 2012, [3]Krysan et al. 1996, [4]Green and Tobin 1999.
*Primer pairs compatible with *A. halleri* (Lan3.1) in addition to *A. thaliana* (Col-0).
[a]Using the published guide for accessing *A. thaliana* T-DNA insertional mutants by O'Malley et al. 2015, and based on the expected orientation and configuration (head-to-head, resulting in a 34-kb insertion in the *CCA1* gene) of the T-DNA in the *cca1-1* mutant as characterized by Green and Tobin 1999, three independent PCR genotyping runs were conducted. These used the LB-*cca1-1* primer in combination with either (1) FP-*cca1-1* (amplicon size = *ca.* 1,000 bp) or (2) RP-*cca1-1* (amplicon size = *ca.* 2,000 bp). PCR products were successfully amplified from genomic DNA of *cca1-1* mutant plants in both reactions, but not in wild-type *A. thaliana* (Col-0) plants. A third reaction using the FP-*cca1-1* and RP-*cca1-1* primers yielded a product only in wild-type plants, but not in mutant plants (amplicon size = *ca.* 3,000 bp).
[b]For genotyping of the construct carrying *GUS* in the crosses with the *cca1-1* mutant allele.
Except where indicated otherwise, the primers listed in this table were generated in this study. The underlined segments in the primer sequences listed above correspond to the nucleotides that hybridize to a target *HMA4* promoter region to be amplified. Bold, non-underlined fonts in italics indicate the nucleotide changes made by site-directed mutagenesis. The primers used for directional cloning (see Methods section) include the four nucleotides, *CACC*, highlighted in italics.

**Supplementary Table S2. Cloning procedures, primers, and plasmids used in this study**

| Construct name, abbreviation | Construct primer pairs | Template | Cloning procedure, PCR cycling parameters | Purpose |
|---|---|---|---|---|
| *AtHMA4*$_P$FL, *At*$_P$FL | [1] | *A. thaliana*$^{\text{Col-0}}$ gDNA | [1] | Generation of a full-length *At*$_P$ |
| *AhHMA4-1*$_P$FL, *Ah-1*$_P$FL | [1] | *A. halleri*$^{\text{Lan 3.1}}$ gDNA | [1] | Generation of a full-length *Ah-1*$_P$ |
| *AhHMA4-2*$_P$FL, *Ah-2*$_P$FL | [1] | *A. halleri*$^{\text{Lan 3.1}}$ gDNA | [1] | Generation of a full-length *Ah-2*$_P$ |
| *AhHMA4-3*$_P$FL, *Ah-3*$_P$FL | [1] | *A. halleri*$^{\text{Lan 3.1}}$ gDNA | [1] | Generation of a full-length *Ah-3*$_P$ |
| *AtHMA4*$_P$ Δ597, Δ597 or *At*$_P$ ΔDR | FP#1+RP#1 | *At*$_P$FL | 1* | Deletion of the distal region in *At*$_P$ |
| *AtHMA4*$_P$ Δ405, Δ405 | FP#2+RP#2 | *At*$_P$FL | 1* | Deletion of part of the intermediate region in *At*$_P$ |
| *AtHMA4*$_P$ Δ332, Δ332 | FP#3+RP#3 | *At*$_P$FL | 1* | Deletion of part of the intermediate region in *At*$_P$ |
| *AtHMA4*$_P$ Δ234, Δ234 | FP#4+RP#4 | *At*$_P$FL | 1* | Deletion of part of the intermediate region in *At*$_P$ |
| *AtHMA4*$_P$ Δ129, Δ129 | FP#5+RP#5 | *At*$_P$FL | 1* | Deletion of the intermediate region in *At*$_P$ |
| *AtHMA4*$_P$ FL Δintron, *At*$_P$FL Δi | FP#6+RP#6 | *At*$_P$FL | 1* | Deletion of the first intron in *At*$_P$ |
| *AhHMA4-1*$_P$ Δ698, Δ698 or *Ah-1*$_P$ ΔDR | FP#7+RP#7 | *Ah-1*$_P$FL | 1* | Deletion of the distal region in *Ah-1*$_P$ |
| *AhHMA4-1*$_P$ Δ438, Δ438 | FP#8+RP#8 | *Ah-1*$_P$FL | 1* | Deletion of part of the intermediate region in *Ah-1*$_P$ |
| *AhHMA4-1*$_P$ Δ327, Δ327 | FP#3+RP#3 | *Ah-1*$_P$FL | 1* | Deletion of part of the intermediate region in *Ah-1*$_P$ |
| *AhHMA4-1*$_P$ Δ230, Δ230 | FP#4+RP#4 | *Ah-1*$_P$FL | 1* | Deletion of part of the intermediate region in *Ah-1*$_P$ |
| *AhHMA4-1*$_P$ Δ130, Δ130 | FP#9+RP#9 | *Ah-1*$_P$FL | 1* | Deletion of the intermediate region in *Ah-1*$_P$ |
| *AhHMA4-1*$_P$FL Δintron, *Ah-1*$_P$FL Δi | FP#10+RP#10 | *Ah-1*$_P$FL | 1* | Deletion of the first intron in *Ah-1*$_P$ |
| *AhHMA4-2*$_P$ Δ515, *Ah-2*$_P$ ΔDR | FP#11+RP#11 | *Ah-2*$_P$FL | 1* | Deletion of the distal region in *Ah-2*$_P$ |
| *AhHMA4-2*$_P$FL Δintron, *Ah-2*$_P$FL Δi | FP#12 RP#12 | *Ah-2*$_P$FL | 1* | Deletion of the first intron in *Ah-2*$_P$ |
| *AhHMA4-2*$_P$ Δ130Δi, *Ah-2*$_P$ ΔDIR | FP#13+RP#13 | *Ah-2*$_P$FL Δi | 1* | Deletion of the intermediate region in *Ah-2*$_P$ |
| *AhHMA4-2*$_P$ Δ61Δi, *Ah-2*$_P$ ΔDIPR | FP#14+RP#14 FP#15+RP#15 | *Ah-2*$_P$FL Δi | 2* | Deletion of the proximal region in *Ah-2*$_P$ |
| *AhHMA4-3*$_P$ Δ310, *Ah-3*$_P$ ΔDR | FP#16+RP#16 | Digested pBKS | 2* | Deletion of the distal region in *Ah-3*$_P$ |
| *AhHMA4-3*$_P$FL Δintron, *Ah-3*$_P$FL Δi | FP#12+RP#12 | *Ah-3*$_P$FL | 2* | Deletion of the first intron in *Ah-3*$_P$ |
| *AhHMA4-3*$_P$ Δ120Δi, *Ah-3*$_P$ ΔDIR | FP#17+RP#14 FP#+RP#15 | *Ah-3*$_P$FL Δi | 2* | Deletion of the intermediate region in *Ah-3*$_P$ |
| *AhHMA4-3*$_P$ Δ82Δi, *Ah-3*$_P$ ΔDIPR | FP#19+RP#16 FP#20+RP#15 | *Ah-3*$_P$FL Δi | 2* | Deletion of the proximal region in *Ah-3*$_P$ |
| *AtHMA4*$_P$ Δ754Δintron, Δ754Δi | FP#21+RP#17 | *At*$_P$FL Δi | 3[a] 66°C[A] - 30 s 72°C - 40 s[E] | Intronless *At*$_P$ construct for introducing the *Ah-1*$_P$ *cis*-elements MHE1a,b and MHE2a,b |
| *AtHMA4*$_P$ Δ597Δi, Δ597Δi or ΔDRΔi | FP#22+RP#17 | *At*$_P$FL Δi | 3[a] 64°C[A] - 30 s 72°C - 30 s[E] | Deletion of the distal region and the first intron of *At*$_P$, and for introducing MHE2a,b |
| Intermediate *AtHMA4*$_P$ Δ754Δi >MHE1b | FP#23+RP#18 | Δ754Δintr | 3[c] 63°C[A] - 30 s 72°C - 40s[E] | Intermediate *At*$_P$ Δ754Δi construct harboring the MHE1b *cis*-regulatory mutations |
| *AtHMA4*$_P$ Δ754Δi >MHE1a,b | FP#24+RP#19 | Intermediate *AtHMA4*$_P$ Δ754Δi >MHE1b | 3[c] 63°C[A] - 30 s 72°C - 40s[E] | Final *At*$_P$ Δ754Δi construct harboring the MHE1a,b *cis*-regulatory mutations |
| *AtHMA4*$_P$ Δ754Δi >MHE2a,b | FP#25+RP#20 | Δ754Δintr | 3[c] 64°C[A] - 30 s 72°C - 40s[E] | *At*$_P$ Δ754Δi construct for introducing the *Ah-1*$_P$ *cis*-elements MHE2a,b |
| *AtHMA4*$_P$ Δ754Δi >MHE1a,b & >MHE2a,b, *At*$_P$ Δ754Δi >MHE1&2 | FP#25+RP#20 | *AtHMA4*$_P$ Δ754Δi >MHE1a,b | 3[c] 64°C[A] - 30 s 72°C - 40s[E] | *At*$_P$ Δ754Δi construct harboring both MHE1a,b and MHE2a,b *cis*-regulatory mutations |
| *AtHMA4*$_P$ Δ129Δintron, Δ129Δi or *At*$_P$ΔDIRΔi | FP#26+RP#17 | *At*$_P$FL Δi | 3[a] 52°C[A] - 30 s 72°C - 15 s[E] | Deletion of the intermediate region and the first intron in *At*$_P$ |
| *AtHMA4*$_P$ Δ88Δintron, Δ88Δi or *At*$_P$ΔDIPRΔi | FP#27+RP#17 | *At*$_P$FL Δi | 3[a] 51°C[A] - 30 s 72°C - 15 s[E] | Deletion of the proximal region and the first intron in *At*$_P$ |
| *AhHMA4-1*$_P$ Δ1,362, Δ1,362 | FP#28+RP#21 | *Ah-1*$_P$FL | 3[a] 58°C[A] - 30 s | Dissection of the distal region in *Ah-1*$_P$ |

| | | | | |
|---|---|---|---|---|
| | | | 72°C - 1:10 min[E] | |
| *AhHMA4-1*$_P$ Δ909, Δ909 | FP#29+RP#21 | Δ1,362 | 3[a] 62°C[A] - 30 s 72°C - 1:00 min[E] | Dissection of the distal region in *Ah-1*$_P$ |
| *AhHMA4-1*$_P$ Δ805, Δ805 | FP#30+RP#21 | Δ909 | 3[a] 62°C[A] - 30 s 72°C – 50 s[E] | Dissection of the distal region in *Ah-1*$_P$ |
| *AhHMA4-1*$_P$ Δ86, Δ86 | FP#31+RP#21 | Δ805 | 3[a] 67°C[A] - 30 s 72°C – 30 s[E] | Control construct to compare promoter activity driven by a basal *Ah-1*$_P$ and *Ah-1*$_P$ constructs in which MHE1&2 has been mutagenized (see *Ah-1*$_P$ FL MHE1/2$_{mut}$ below) |
| Intermediate AB fragment | FP#32+RP#22 | *Ah-1*$_P$FL | 3[b] 62°C[A] - 30 s 72°C – 45 s[E] | Intermediate AB fragment including the distal and intermediate region of *Ah-1*$_P$ |
| Intermediate CD fragment | FP#33+RP#21 | *Ah-1*$_P$FL | 3[b] 55°C[A] - 30 s 72°C – 30 s[E] | Intermediate CD fragment, including the entire downstream region (5'UTR, intron, and 30 bp CDS fragment) of the proximal region of *Ah-1*$_P$ |
| *AhHMA4-1*$_P$ FL ΔER2, ΔER2 | FP#34+RP#21 | AB+CD fragments** | 3[b] 55°C[A] - 30 s 72°C – 1:30 min[E] | Final construct in which the enhancing, proximal, region in *Ah-1*$_P$ has been deleted |
| *AhHMA4-1*$_P$ Δ130Δintron, ΔDIRΔi | FP#35+RP#21 | *Ah-1*$_P$FL Δi | 3[a] 56°C[A] - 30 s 72°C – 15 s[E] | Deletion of the intermediate region and the first intron in *Ah-1*$_P$ |
| *AhHMA4-1*$_P$ Δ86Δintron, ΔDIPRΔi | FP#36+RP#21 | *Ah-1*$_P$FL Δi | 3[a] 67°C[A] - 30 s 72°C – 15 s[E] | Deletion of the proximal region and the first intron in *Ah-1*$_P$ |
| Intermediate *AhHMA4-1*$_P$ FL MHE1b$_{mut}$ | FP#37+RP#23 | *Ah-1*$_P$FL | 3[c] 60°C[A] - 30 s 72°C – 1:15 s[E] | Intermediate *AhHMA4-1*$_P$ FL MHE1b$_{mut}$ construct for site-directed mutagenesis of MHE1a,b |
| *AhHMA4-1*$_P$FL MHE1$_{mut,}$ | FP#38+RP#24 | Intermediate *AhHMA4-1*$_P$FL MHE1b$_{mut}$ | 3[c] 65°C[A] - 30 s 72°C – 1:15 s[E] | Final *AhHMA4-1*$_P$ FL MHE1b$_{mut}$ construct for site-directed mutagenesis of both MHE1a and b |
| *AhHMA4-1*$_P$FL MHE2$_{mut}$ | FP#39+RP#25 | *Ah-1*$_P$FL | 3[c] 63°C[A] - 30 s 72°C – 1:15 s[E] | *AhHMA4-1*$_P$ FL construct for site-directed mutagenesis of both MHE2a and b |
| *AhHMA4-1*$_P$ FL MHE1/2$_{mut}$ | FP#39+RP#25 | *AhHMA4-1*$_P$FL MHE1$_{mut}$ | 3[c] 63°C[A] - 30 s 72°C – 1:15 s[E] | Final *AhHMA4-1*$_P$ FL MHE1a,b$_{mut}$ construct for site-directed mutagenesis of both MHE2a and b |
| *AtHMA4*$_P$ FL Δcds | FP#40+RP#26 | *At*$_P$FL | 3[c] 61°C[A] - 30 s 72°C – 1:40 s[E] | Full-length *AtHMA4*$_P$ lacking the 5' 204 bp coding sequence |
| Intermediate AB fragment | FP#41+RP#27 | *Ah-1*$_P$FL | 3[b] 60°C[A] - 30 s 72°C – 15 s[E] | Intermediate AB fragment including the distal region of *Ah-1*$_P$ |
| Intermediate CD fragment | FP#42+RP#28 | *At*$_P$FL | 3[b] 65°C[A] - 30 s 72°C – 15 s[E] | Intermediate CD fragment including the downstream region of *At*$_P$, starting at IR and until the cds |
| *AhHMA4-1*$_P$DR:*AtHMA4*$_P$IR-cdsΔintron, *Ah-1*$_P$DR:*At*$_P$IR-cdsΔi, | FP#43+RP#28 | AB+CD fragments** | 3[b] 66°C[A] - 30 s 72°C – 1:00 min[E] | Final swap construct in which the DR of *Ah-1*$_P$ has fused to the downstream region of *At*$_P$, starting at IR and until the cds |

[1]Hanikenne et al. 2008.
*PCR cycling conditions are given in Materials and Methods.
[a-c]PCR-mediated amplification, overlap PCR extension, and quick change site-directed mutagenesis respectively — see Methods section.
[A]Annealing temperature.
[E]Extension time.
**AB and CD fragments were used in equal quantities of each as described by Heckman and Pease 2007.

**Supplementary Table S3. Motifs identified in ER1$^+$ and ER2 segments of *AhHMA4* promoters of *A. halleri* accessions**

| Motif no. and sequence (5’-3’) | *e*-value | Width (nt) |
|---|---|---|
| Enhancing Region 1 (ER1$^+$) | | |
| 1. CYWTGTAACCWT | 1.4*e*-355 | 12 |
| 2. TTTTTWTTWATW | 8.2*e*-125 | 12 |
| 3. TTTAAATTA | 9.8*e*-006 | 9 |
| 4. WWAWW | 1.5*e*+007 | 5 |
| Enhancing Region 2 (ER2) | | |
| 1. MAMTATCT | 4.8*e*-688 | 8 |
| 2. TTACAAAAGAA | 2.2*e*-220 | 11 |
| 3. ATAATATCTAA | 1.1*e*-059 | 11 |

Alphabet symbols correspond to the annotation implemented in the Tomtom tool within the MEME suite and are defined as follows: Y : pyrimidine (C or T), W : weak bond (A or T), M : amino bases (A or C).
